# Multivariate genetic analysis of personality and cognitive traits reveals abundant pleiotropy and improves prediction

**DOI:** 10.1101/2022.02.28.481967

**Authors:** Guy Hindley, Alexey Shadrin, Dennis van der Meer, Nadine Parker, Weiqiu Cheng, Kevin S. O’Connell, Shahram Bahrami, Aihua Lin, Naz Karadag, Børge Holen, Thomas Bjella, Chun C Fan, Torill Ueland, Srdjan Djurovic, Olav B. Smeland, Oleksandr Frei, Anders M. Dale, Ole A. Andreassen

**Affiliations:** NORMENT Centre, Institute of Clinical Medicine, University of Oslo and Division of Mental Health and Addiction, Oslo University Hospital, 0407 Oslo, Norway; Psychosis Studies, Institute of Psychiatry, Psychology and Neurosciences, King’s College London, 16 De Crespigny Park, London SE5 8AB, United Kingdom; KG Jebsen Centre for Neurodevelopmental disorders, University of Oslo, Oslo, Norway; School of Mental Health and Neuroscience, Faculty of Health, Medicine and Life Sciences, Maastricht University, Maastricht, The Netherlands; Division of Mental Health and Addiction, Oslo University Hospital, Oslo, Norway; Department of Cognitive Science, University of California, San Diego, La Jolla, CA, USA; Multimodal Imaging Laboratory, University of California San Diego, La Jolla, CA 92093, USA; Department of Psychology, University of Oslo, Norway; Department of Medical Genetics, Oslo University Hospital, Oslo, Norway; NORMENT Centre, Department of Clinical Science, University of Bergen, Bergen, Norway; Center for Bioinformatics, Department of Informatics, University of Oslo, PO box 1080, Blindern, 0316 Oslo, Norway; Department of Psychiatry, University of California, San Diego, La Jolla, CA, USA; Department of Neurosciences, University of California San Diego, La Jolla, CA 92093, USA; Department of Radiology, University of California, San Diego, La Jolla, CA 92093, USA

**Author notes:** **Corresponding authors** Correspondence: Guy Hindley, Alexey Shadrin Ole A Andreassen. These authors contributed equally (Guy Hindley, Alexey Shadrin).

**Keywords:** Cognition, personality, Big 5 personality, neuroticism, GWAS, multivariate GWAS, MOSTest, pleiotropy

## Abstract

Personality and cognition are heritable mental traits, and their genetic determinants may be distributed across interconnected brain functions. However, previous studies have employed univariate approaches which reduce complex traits to summary measures. We applied the “pleiotropy-informed” multivariate omnibus statistical test (MOSTest) to genome-wide association studies (GWAS) of 35 item and task-level measures of neuroticism and cognition from the UK Biobank (n=336,993). We identified 431 significant genetic loci and found evidence of abundant pleiotropy across personality and cognitive domains. Functional characterisation implicated genes with significant tissue-specific expression in all tested brain tissues and enriched in brain-specific gene-sets. We conditioned independent GWAS of the Big 5 personality traits and cognition on our multivariate findings, which boosted genetic discovery in other personality traits and improved polygenic prediction. These findings advance our understanding of the polygenic architecture of complex mental traits, indicating a prominence of pleiotropic genetic effects across higher-order domains of mental function.

**Graphical abstract:** 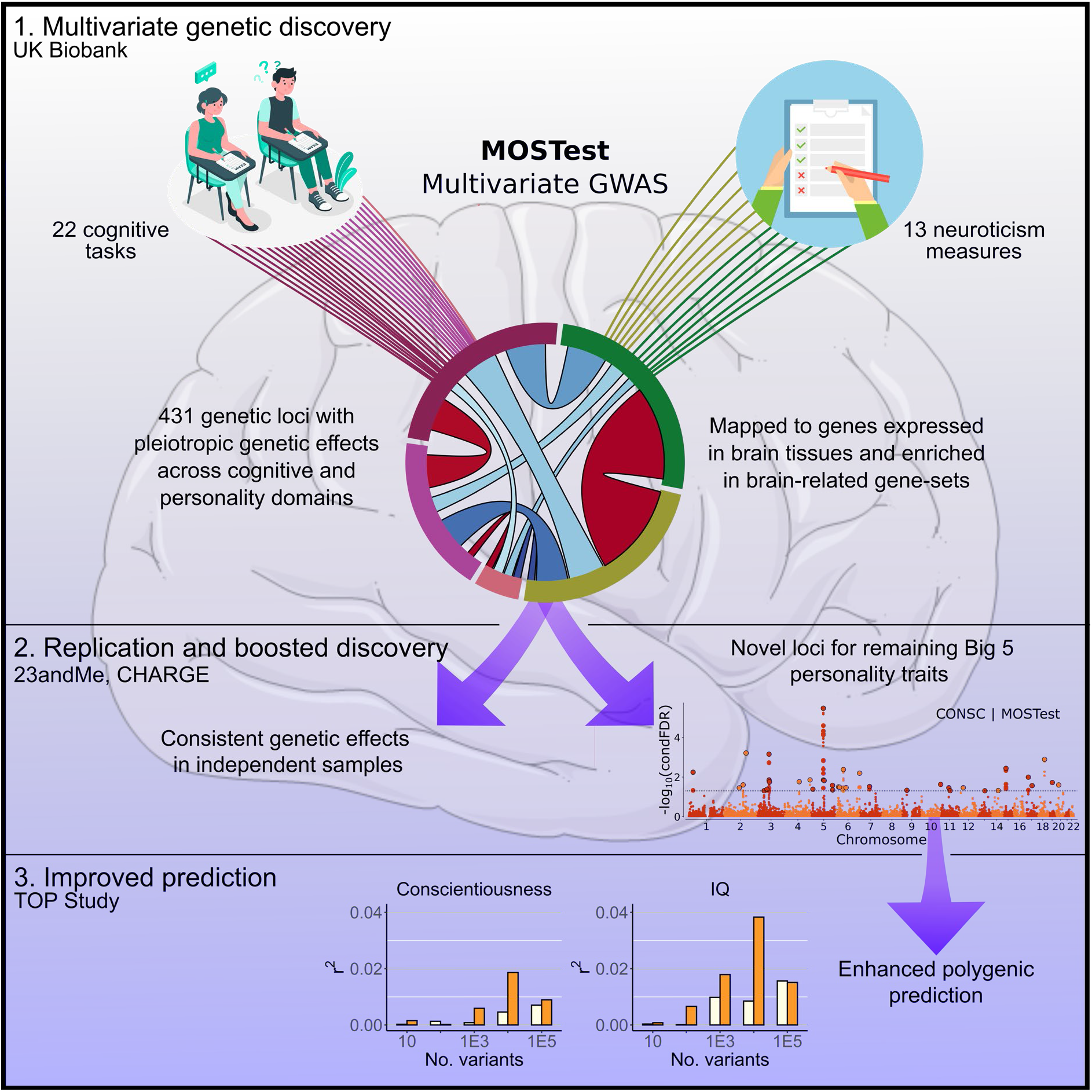

## Introduction

The brain is responsible for a diverse set of interconnected and overlapping functions. Among these, personality and cognition both represent heritable, higher-order domains of mental functioning that (i) remain relatively stable between late adolescence and old age (Damian et al., 2019; Walhovd et al., 2016), (ii) form central components of an individual’s identity, and (iii) are related to multiple physical and mental health outcomes (Strickhouser et al., 2017; Wraw et al., 2015). They are also interrelated, with evidence of complex patterns of association between personality structure, cognitive functioning (Wettstein et al., 2017) and academic performance (Mammadov, 2021). A comprehensive investigation of their genetic foundations can provide insights into the neurobiological mechanisms influencing these fundamental human traits.

Accelerated by the population-based cohort the UK Biobank (UKB; n=~500,000), genome-wide association studies (GWAS) have revealed evidence of genetic overlap between personality and cognitive traits. Thirty-eight genetic loci were shared between 136 loci associated with neuroticism (Nagel et al., 2018a), one of the “Big 5” personality traits defined as the propensity to experience negative emotions (Widiger and Oltmanns, 2017), and 205 loci associated with general intelligence (Savage et al., 2018), defined as the “common factor” underlying diverse cognitive functions. Neuroticism and general intelligence also exhibit weak but significant negative genetic correlation (r_g_=− 0.16) (Widiger and Oltmanns, 2017) and higher polygenic scores (PGS) for neuroticism predict lower intelligence (Hill et al., 2020).

Both GWAS described employ univariate analytical approaches, which reduce complex mental traits to a single measure (Nagel et al., 2018a; Savage et al., 2018). The limitations of this approach are underscored by an item-level analysis of neuroticism which found that, despite negative genetic correlation at the sum-score level, two neuroticism sub-factors were positively genetically correlated with general intelligence (Hill et al., 2020). A second item-level analysis of the neuroticism scale also showed divergent patterns of genetic correlation between individual neuroticism items and diverse mental traits (Nagel et al., 2018b).

In contrast, multivariate approaches simultaneously model the matrix of correlations between phenotypes, thus more accurately representing the interconnected nature of the brain and its functions. Multivariate analysis can also increase statistical power in mental traits, as demonstrated by a study of neuroticism items in UKB which used canonical correlation analysis (CCA) to discover twice the number of genetic loci compared to univariate GWAS (Nagel et al., 2018b). A boost in genetic discovery has also been demonstrated by the “pleiotropy-informed” multivariate omnibus statistical test (MOSTest). Applying MOSTest to brain imaging phenotypes has shown that alterations in brain morphology and functional connectivity are associated with hundreds of genetic loci with “pleiotropic” genetic effects across the brain, even despite weak genetic correlation (van der Meer et al., 2020a, 2021; Roelfs et al., 2022; Shadrin et al., 2021). We hypothesised that the genetic architecture of interconnected higher-order mental traits, such as cognition and personality are driven by similar pleiotropic effects.

Our understanding of the genetics of personality traits beyond neuroticism is limited, in part because UKB did not collect data on the four remaining personality traits within the “Big 5” taxonomy. As such, only eight loci have been reported across all five measures in the largest GWAS to date (n=76,600-122,886) (Lo et al., 2017). However, it is possible to boost statistical power for genetic discovery, identify shared genetic loci and improve prediction in underpowered GWAS by leveraging genetic overlap with a second, more powerful GWAS using the conditional false discovery rate framework (cFDR) (Andreassen et al., 2013; van der Meer et al., 2020b; Smeland et al., 2019a). This approach has recently been applied to MOSTest analyses of brain structural (van der Meer et al., 2020b) and functional measures (Roelfs et al., 2022) to improve discovery and prediction of mental disorders.

Given evidence of genetic overlap between neuroticism and cognition, we sought to boost the statistical power for genetic discovery by exploiting pleiotropic genetic effects across item and task-level measures of neuroticism and cognition. By applying “pleiotropy-informed” MOSTest, which incorporates scenarios of mixed effect directions, we found a substantial boost in discovery driven by shared genetic effects across domains. The widespread effects were supported by functional analysis, which identified underlying neurobiological processes distributed across brain regions. We additionally leveraged our multivariate analysis to boost genetic discovery across the remaining Big Five personality traits and improve polygenic prediction.

## Results

### Sample description

The UKB is a population-based cohort comprising over 500,000 participants between the ages of 39-72 (Bycroft et al., 2018). At enrolment, all participants were invited to complete a touchscreen questionnaire, including 12 dichotomous items derived from the neuroticism subscale of the Eysenck Personality Questionnaire-Revised Short Form (Eysenck et al., 1985). They additionally completed 25 diverse cognitive tasks, either at enrolment or during follow-up visits. These included measures of fluid intelligence, reaction time, executive function, and memory (Cullen et al., 2017) (***table 1, supplementary table 1***). After also calculating sum-scores for neuroticism and fluid intelligence, we included all 39 measures to maximise statistical power for genetic discovery. After removing participants of non-White European ancestry and related individuals, the mean sample size across all 35 measures was 214,974, ranging from 11,679 to 336,993 (***table 1, supplementary figure 1***). Sample sizes were more variable among cognitive tasks than neuroticism items. Mean age was 56.9 years (s.d.=8.0) at enrolment and 53.7% of included participants were female.

**Table 1:**
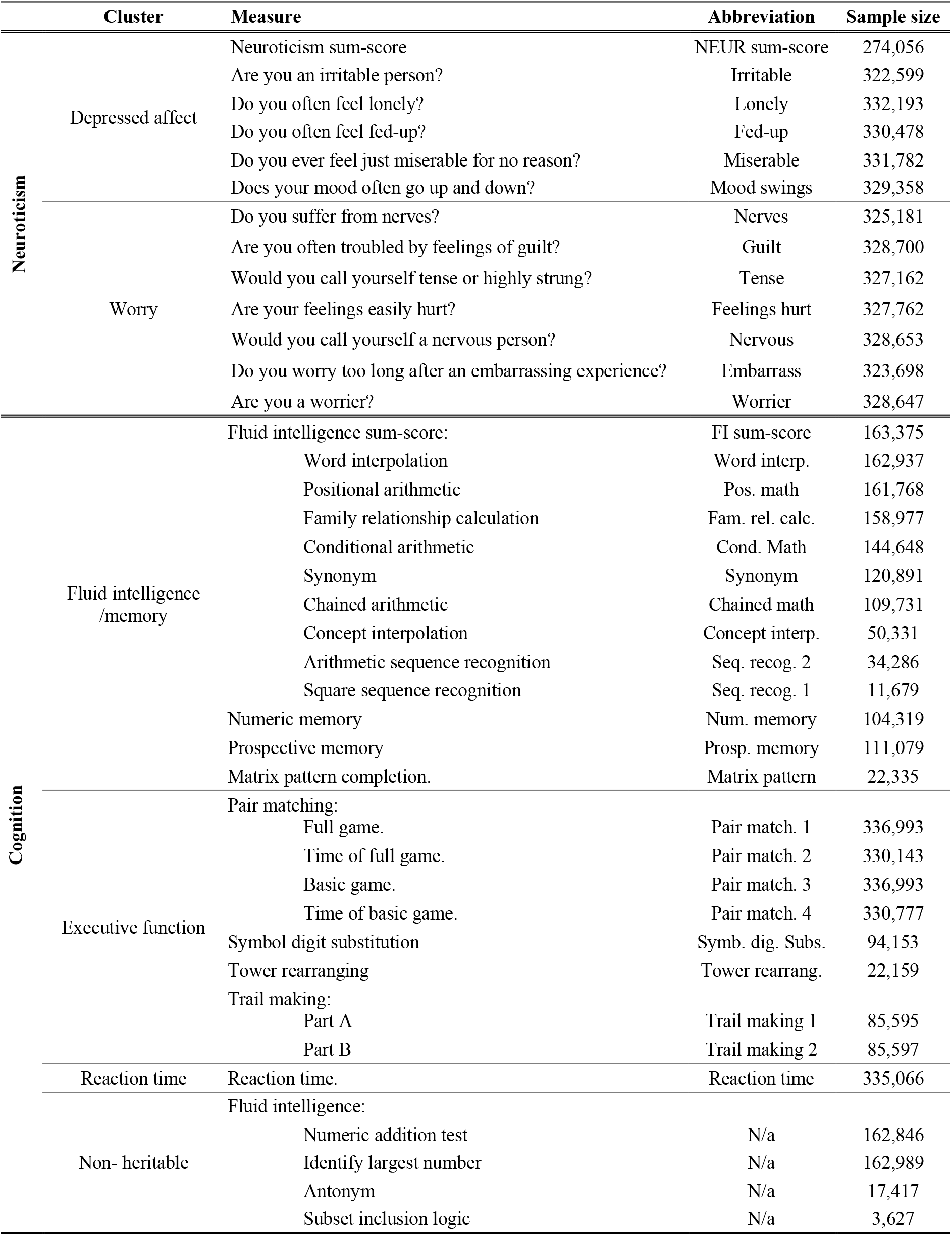
Overview of neuroticism and cognitive measures from UK Biobank. Clusters are derived from genetic correlation-based hierarchical clustering (*figure 1*). Further details are provided in *supplementary table 1*.

### Item-level heritability and genetic correlations

To provide an overview of the heritability of item and task-level measures, we first calculated linkage-disequilibrium score regression (LDSR) SNP-heritabilities (h^2^_SNP_) for all included measures (***supplementary figure 1, supplementary table 2***). Mean h^2^_SNP_ for all 39 items was 0.083 (s.d.=0.052). The neuroticism sum-score (h^2^_SNP_=0.12, s.e.=0.0053, p=2.62e-^106^) and fluid intelligence sum-score (h^2^_SNP_=0.21, s.e.=0.008, p=2.54e-^154^) were the most heritable measures within each domain, respectively. Four conditions within the fluid intelligence scale were not significantly heritable. Among these, “numeric addition test” and “identify largest number” displayed highly skewed responses, most likely due to the simplicity of the tasks. In contrast the conditions “antonym” and “subset inclusion logic” were underpowered (n=3,627-11,679) as they were performed at the end of a timed session. Since the inclusion of non-heritable phenotypes may reduce statistical power (van der Meer et al., 2020a), these four measures were removed from the rest of the analysis, leaving a total of 35 measures.

We next computed genetic correlations using LDSR for all pairs of measures followed by hierarchical clustering to explore directional genetic relationships across included measures (***figure 1, supplementary results***) (Bulik-Sullivan et al., 2015a). Neuroticism and cognitive measures shared weak negative genetic correlations (mean r_g_=−0.15, s.d.=0.12) compared to moderate to strong positive correlations within each domain (neuroticism measures mean r_g_=0.64, s.d.=0.14; cognitive measures mean rg=0.56, s.d.=0.23). Consequently, neuroticism and cognitive measures clustered separately. Neuroticism items further clustered into two sub-groups, mapping onto anxiety related features (“worry”) and depressive features (“depressed affect”), replicating previous findings (Nagel et al., 2018b). Cognitive measures were more heterogenous, with “reaction time” distinct from two larger clusters relating to fluid intelligence, prospective memory, and numeric memory (“fluid intelligence/memory”) and executive function and visuospatial memory (“executive function”). A similar pattern was observed among phenotypic correlations (***figure 1, supplementary results***).

**Figure 1:**
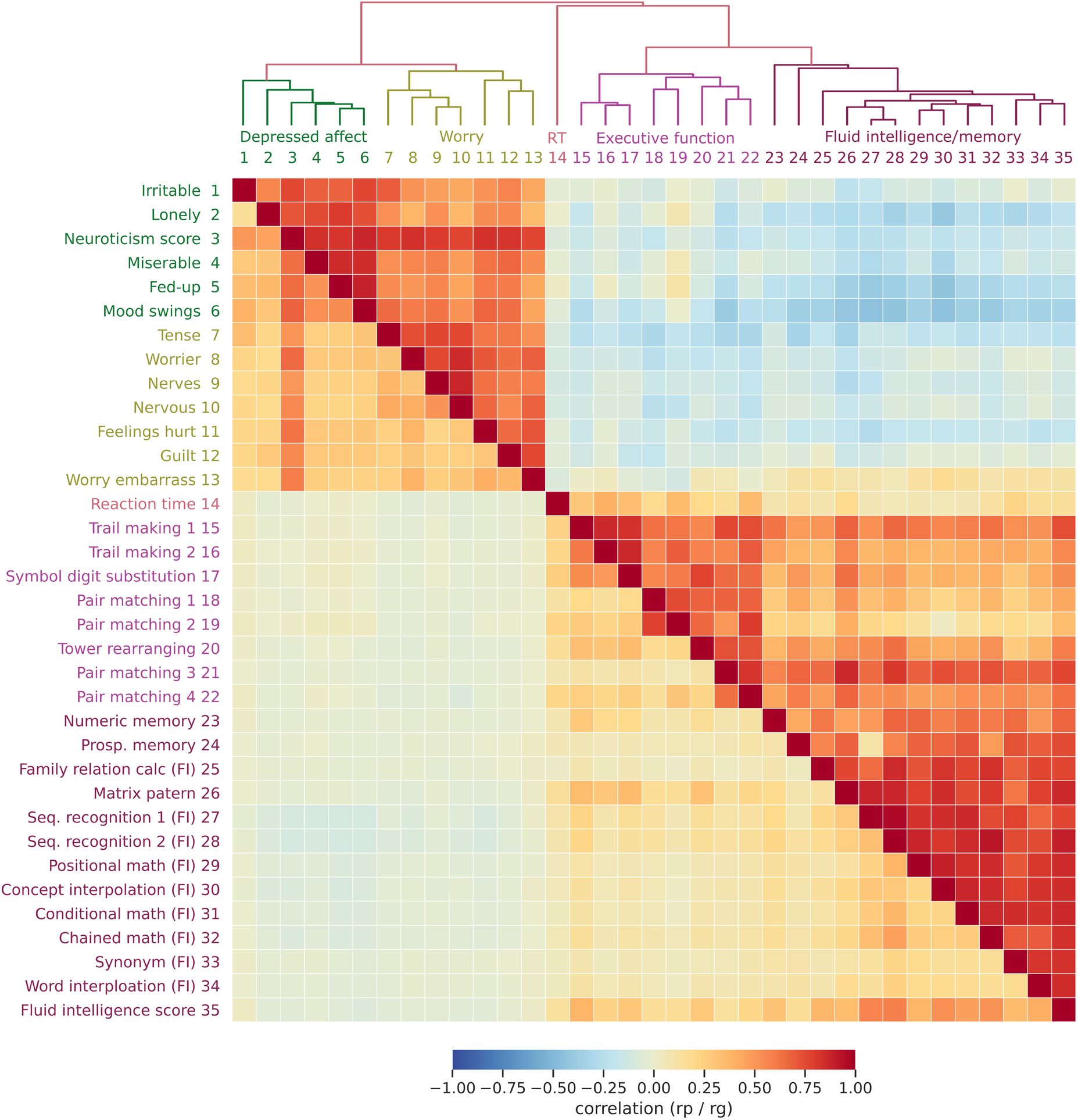
Heatmap of genetic and phenotypic correlations across mental traits. LDSR genetic correlations (rg, top right) and Spearman rank phenotypic correlations (rp, bottom left) reveal a pattern of moderate to strong positive genetic correlations within neuroticism and cognitive domains but weak negative genetic correlations across cognition and neuroticism measures. Phenotypically, there were also stronger positive correlations within domains but minimal correlation across domains. Measures were clustered on genetic correlation, revealing 2 neuroticism clusters aligning with previously reported clusters “depressed affect” and “worry” (Nagel et al., 2018b), and 3 cognition clusters, broadly mapping on to “reaction time”, “executive function” and “fluid intelligence/memory”.

### Multivariate GWAS identifies 431 genetic loci with pleiotropic genetic effects

On application of MOSTest to discover pleiotropic genetic effects, we identified 431 independent genetic loci significantly associated with the multivariate distribution of the 35 measures of neuroticism and cognition. This represented a 3.8x boost in locus discovery compared to mass univariate GWAS with correction for multiple testing (“min-P”), which identified 113 loci (***figure 2a, supplementary figure 2, supplementary tables 3-4***). Since MOSTest specifically leverages pleiotropy, this boost in discovery supports the hypothesis of pleiotropic genetic effects across mental traits. We also performed MOSTest analyses on neuroticism and cognitive measures separately to test how much of the boost in locus discovery was driven by cross-domain pleiotropy. Cognitive measures were associated with 221 genetic loci and neuroticism measures were associated with 199 loci. However, 153 loci discovered by the combined MOSTest analyses were not identified by either of the separate analyses, indicating that 35% of the discovered loci were driven by cross-domain pleiotropy.

**Figure 2:**
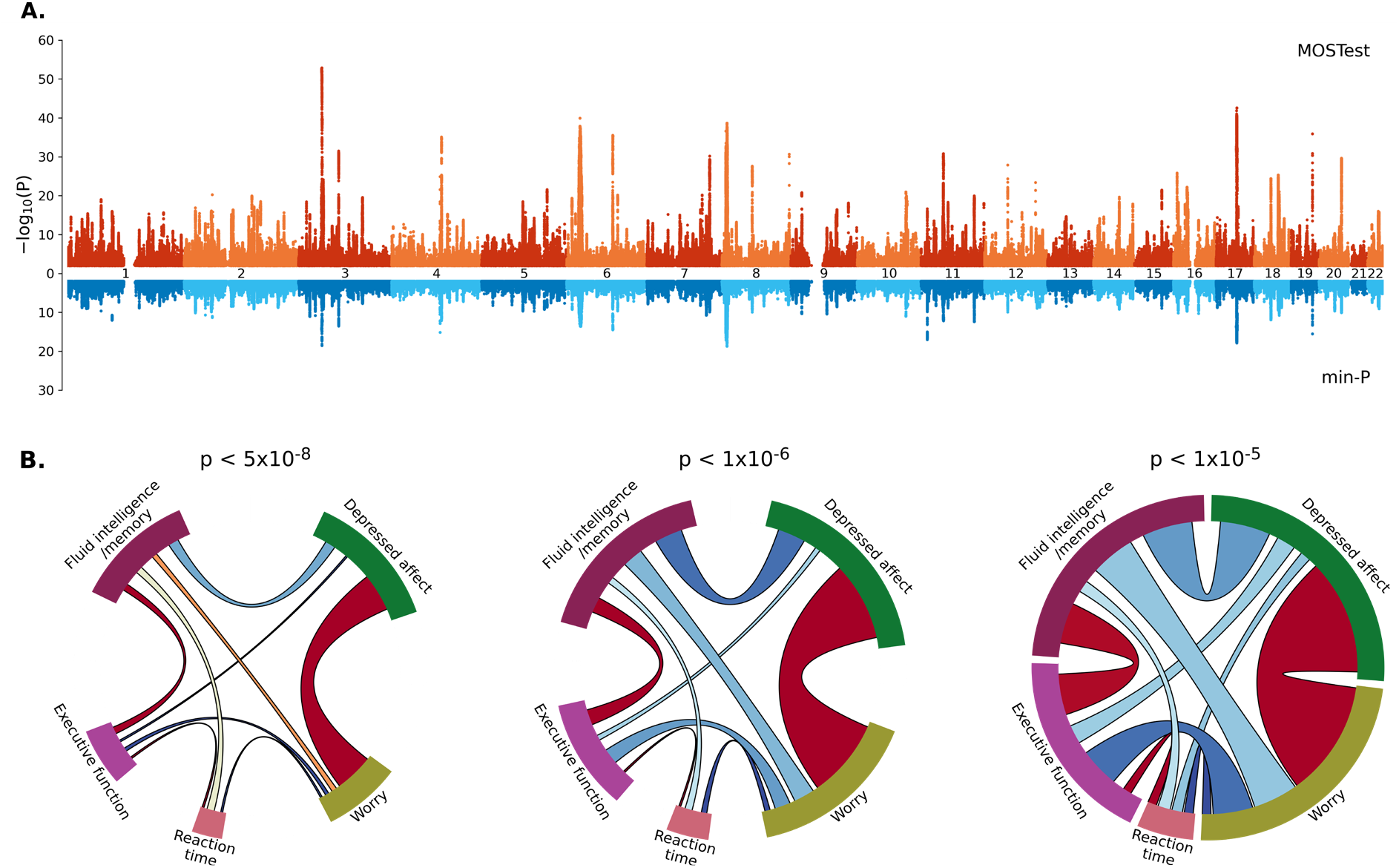
Boosting the signal of genetic association for 35 mental traits by leveraging pleiotropy. **A**. Miami plot for MOSTest (orange) and min-P (blue), plotting each SNP’s −log10(p-values) against chromosomal position. By applying a multivariate framework which leverages pleiotropic effects, there is a substantial boost in signal compared to a “mass univariate” approach such as min-P, evidenced by smaller p-values and a larger number of discovered loci (n=431 vs. 113). This indicates the presence of pleiotropic genetic effects across mental traits. **B**. Shared genetic associations of lead variants across 5 genetic correlation-based clusters (***figure 1***) at three significance thresholds. The number of lead variants within each cluster individually at each significance threshold is represented by the size of the coloured segments. The number of lead variants shared between each pair of clusters is represented by the width of the coloured ribbons. The proportion of variants with concordant effect directions on each cluster is represented by the colour of the ribbons from blue (0) to red (1).

To further illustrate the distribution of genetic effects, we tested for cross-cluster genetic overlap among the 431 lead variants irrespective of effect direction using univariate GWAS p-values from each included measure (***figure 2b, supplementary table 5***). This showed that there was an increase in the number of shared variants at decreasing significance thresholds (p<5×10^−8^, p<1×10^−6^, p<1×10^−5^), indicating that the pleiotropic genetic variants captured by MOSTest had predominantly weak, sub-threshold associations. When comparing across clusters, the two neuroticism clusters “depressed affect” and “worry” shared the largest number of lead variants at all thresholds (n=22-68). Nonetheless, there was a comparable number of shared variants between cognitive and neuroticism clusters (n=0-29) and within cognitive clusters (n=1-24). Although these findings are partly affected by differences in sample size across measures, this provides further evidence of pleiotropic genetic effects across mental traits. We also provide evidence of gene-level overlap across clusters (***supplementary results, supplementary figure 3***).

We compared effect directions of shared lead variants across each pair of clusters at different significance thresholds and calculated the proportion of variants with concordant effect directions on each pair of traits (***figure 2b, supplementary table 5***). This showed that lead variants shared between neuroticism clusters, and between “fluid intelligence/memory” and “executive function”, and “reaction time” and “executive function” possessed highly concordant effects at all significance thresholds (0.98-1.00 concordance), consistent with the strong positive genetic correlations observed in ***figure 1***. In contrast, there was a predominance of variants with discordant effects between “reaction time” and “fluid intelligence/memory” (0.38-0.50 concordance). When comparing across neuroticism and cognitive domains, most shared variants had discordant effects, although there were more prominent mixed effect directions, with concordance ranging from 0-0.33 across all significance thresholds. This is somewhat consistent with the weak genetic correlations between cognitive and neuroticism measures observed in figure 1, although the predominance of discordant lead variants between “executive function” and “worry” (2/23, 0.08 concordance) and “fluid intelligence/memory” and “depressed affect” clusters (5/29, 0.17 concordance) suggests that, to some extent, discovered variants exhibit more strongly discordant genetic effects than the genome-wide average represented by genetic correlations.

To investigate this further, we performed hierarchical clustering of univariate z-scores from all 431 lead variants (***supplementary figures 4-5***). This revealed that the majority of lead variants had discordant effect directions between neuroticism and cognitive measures. However, a minority of variants had either concordant effects across all measures, or had mixed effects within cognitive and neuroticism clusters. This indicates the presence of mixed genetic effect directions but a predominance of discordant effects across domains.

We plotted univariate GWAS p-values from all 35 measures for the top 40 lead variants to illustrate item and task-level patterns of genetic association (***supplementary figure 6***). Plots for five of these variants are presented in ***figure 3***, each exemplifying a distinct pattern of association. While some variants were genome-wide significant in neuroticism clusters (n=76) (***figure 3c)*** or cognitive clusters only (n=85) (***figure 3d-e***), 10 variants were genome-wide significant in measures across both neuroticism and cognitive clusters (***figure 3a***). Nonetheless, most variants had sub-threshold effects (n=260), demonstrating the boost in power provided by multivariate analysis (***figure 3b***). We also present the effect directions at the individual variant level, showing that 4 of the 5 lead variants exhibit a discordant relationship between neuroticism and cognition, consistent with negative genetic correlations. However, ***figure 3b*** exemplified mixed effect directions, with weak positive effects on both “depressed affect” items and cognitive clusters but negative effects on “worry”.

**Figure 3.**
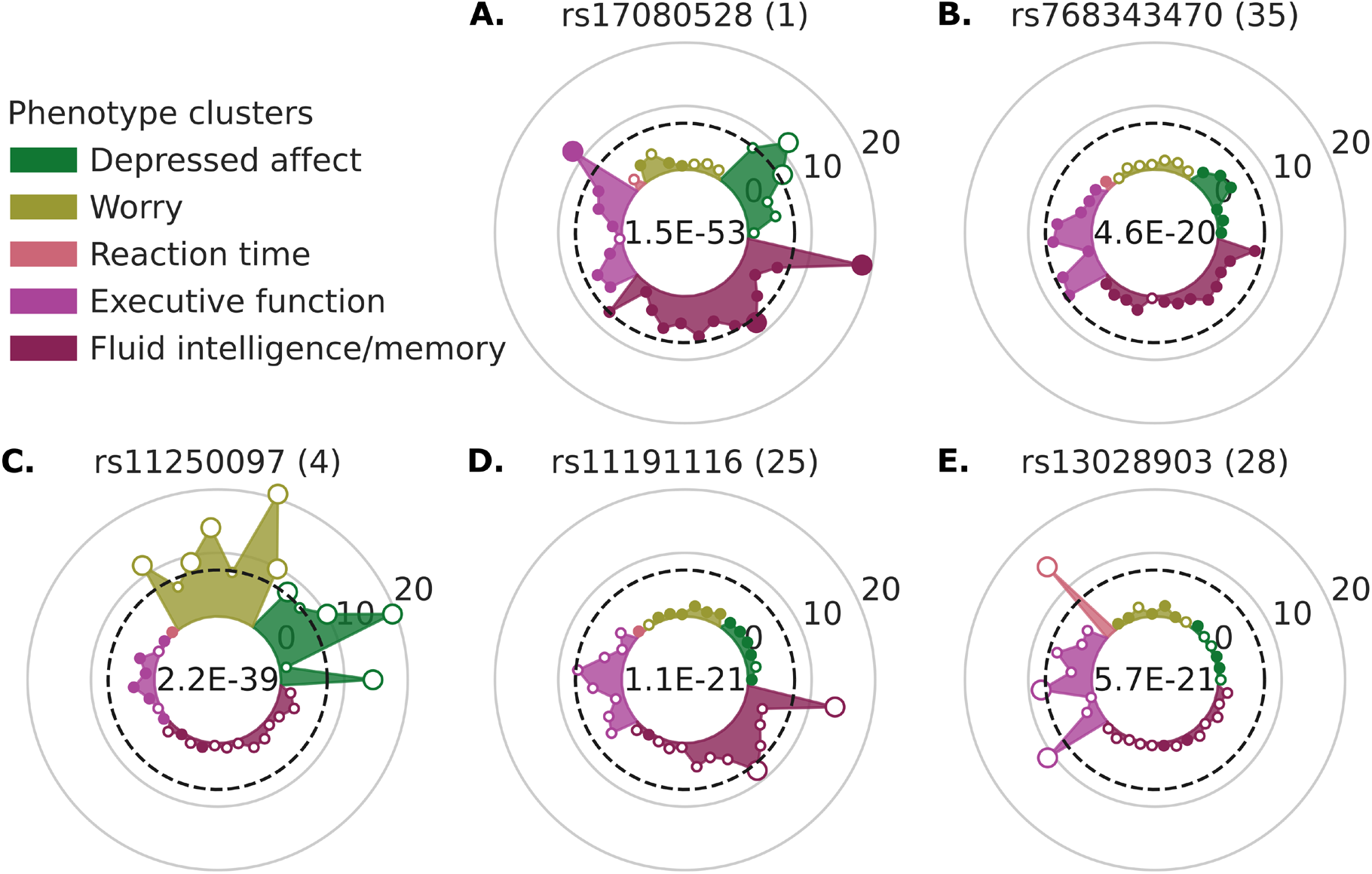
Patterns of pleiotropic genetic associations at the SNP-level. P-values from univariate GWAS of 35 mental traits plotted for 5 lead variants, selected to illustrate five distinct patterns of association. The locus number, which corresponds to MOSTest significance rank, is provided in brackets. Univariate p-values are plotted on the logarithmic scale as the distance from the centre of each circular plot. Genome-wide significance (p<5×10^−8^) is represented by the dashed line. Positive effect direction is illustrated by a filled circle and negative effect direction by a clear circle. Phenotype clusters are derived from genetic correlation-based hierarchical clustering (***figure 2***) **A**: SNP which is genome-wide significant across neuroticism measures (with apparent specificity for the “depressed affect” cluster) and cognitive measures (both in “executive function” and fluid intelligence/memory” clusters), with predominantly positive effects on cognitive tasks and negative effects in neuroticism items. **B**: SNP which is non-significant across all measures, with indication of weak association with “depressed affect” cluster, “executive function” and “fluid intelligence/memory” clusters, and predominantly concordant effects in “cognitive tasks and depressed affect” items. **C**: SNP with genome-wide significance across neuroticism measures but minimal association with cognitive measures, and negative effects on neuroticism items and predominantly positive effects on “executive function”. **D**: SNP with genome-wide significance with “fluid intelligence/memory”, sub-threshold association with “executive function” and minimal association with neuroticism measures, and predominantly negative effects on cognitive tasks and positive effects on neuroticism items. **E**: SNP with genome-wide significance with “reaction time” and “executive function” but minimal association with “fluid intelligence/memory” and neuroticism measures, and negative effects in “fluid intelligence/memory” but weak, mixed effects in all other measures.

### Replication in independent samples

We tested for nominal significance and consistency of effect direction for MOSTest-discovered lead variants in independent samples, including 23andMe neuroticism GWAS (n=59,225) (Lo et al., 2017) and CHARGE “general cognitive function” GWAS (n=53,949) (Davies et al., 2015) (***supplementary table 6***). Out of 140 lead variants which were present in all three samples and had non-ambiguous alleles, 28 were nominally significant in the 23andMe, Inc. neuroticism GWAS, 29 in the CHARGE general cognitive function GWAS, and seven in both datasets. In line with previous studies, we also tested for consistent genetic effects of lead variants across discovery and replication datasets (Lee et al., 2018; Ripke et al., 2020). Since they were the most comparable phenotypes within our analyses, we compared univariate GWAS effect directions for the neuroticism and fluid intelligence sum-scores with 23andMe neuroticism and CHARGE general cognitive function summary statistics, respectively. 102 had concordant effects in neuroticism (exact binomial p=1.61×10^−8^) and 103 had concordant effects in cognition (p=5.67×10^−9^). Seventy-six variants were concordant in both neuroticism and cognition (p=8.57e^−8^), providing additional evidence of pleiotropic effects in independent samples.

### Functional characterisation

Using FUMA (fuma.ctglab.nl) (Watanabe et al., 2017), we performed functional annotation to provide biological insights into the genetic associations captured by MOSTest. We first used multi-marker analysis of genomic annotation (MAGMA) which tests for the association between phenotypic variation and aggregated GWAS p-values for 18,952 human protein-coding genes irrespective of effect direction (de Leeuw et al., 2015). MAGMA identified 1062 multiple comparison-corrected significant genes associated with the 35 measures of neuroticism and cognition (***supplementary table 7***). Next, MAGMA-based tissue specific expression analysis demonstrated highly specific enrichment of mapped genes in brain tissues. At the general tissue level (n=30), the brain, pituitary, ovary and testis were significantly enriched (***supplementary figure 7***). At the detail tissue level (n=53), all of the 14 included brain tissues were significantly enriched, as well as testicular tissue (***figure 4, supplementary figure 8***). When applied to Gene Ontology and canonical pathways there was a clear predominance of brain-related gene-sets. Twenty-nine out of 43 gene-sets were directly implicated in the structure or function of the central nervous system, and eight out of the top 10 significantly enriched gene-sets were related to synaptic structure or function (***figure 4***). Outside of the top 10, other notable gene-sets included “observational learning”, “behavior” and “cognition”, in addition to several neurodevelopmental gene-sets and “gamma aminobutyric acid signalling pathway” (***supplementary table 8***).

**Figure 4.**
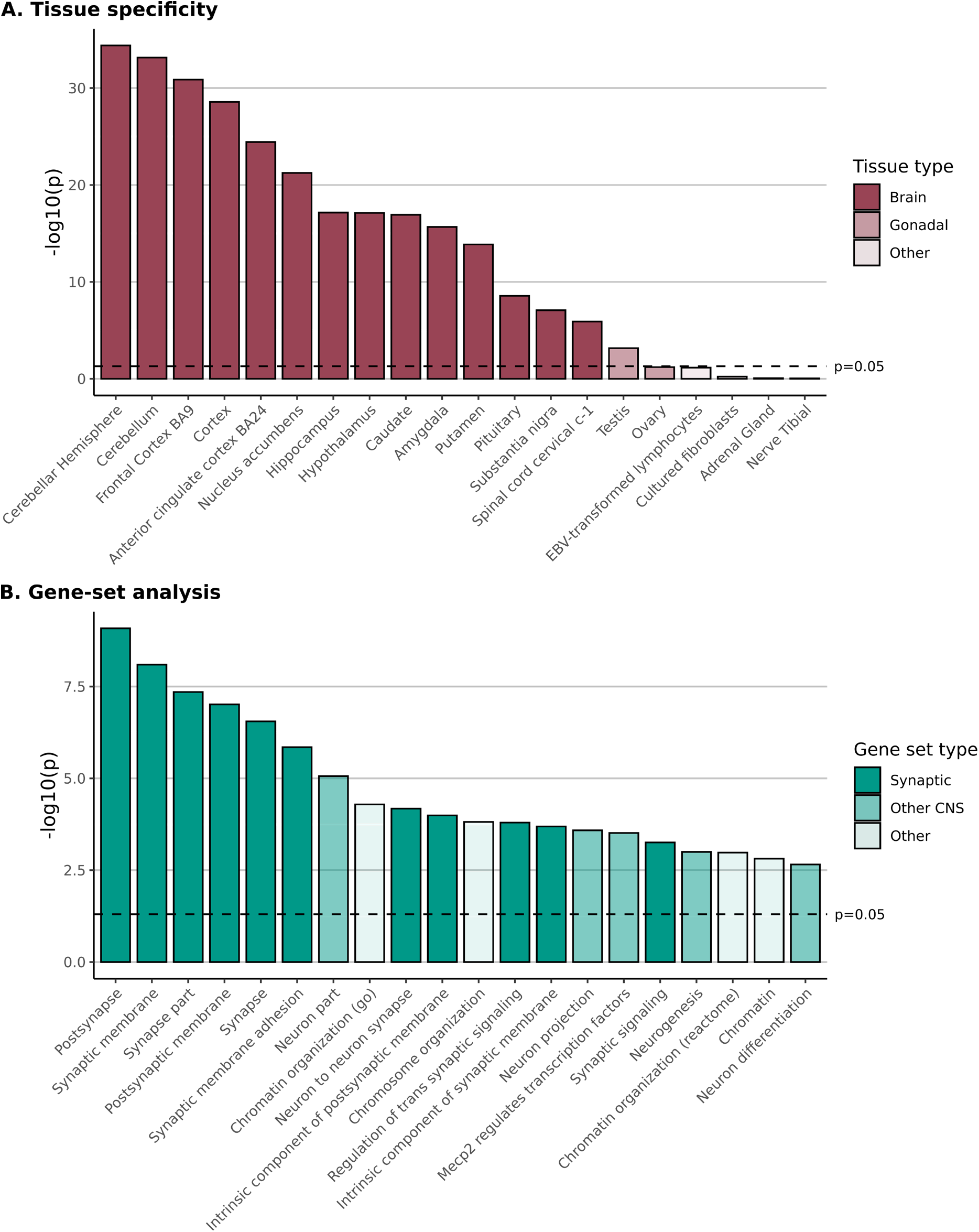
MAGMA tissue specific gene expression and gene-set enrichments. **A**. MAGMA-based tissue specificity analysis of multivariate GWAS of 35 mental traits shows highly specific enrichment across all brain tissues and the testis. All tissues with corrected p<0.1 are presented. All tissues tested are shown in ***supplementary figures 7***. Please note that the ovary was significant when tested at the “general tissues” level (***supplementary figure 8***). **B**. Top 20 gene-sets significantly enriched for gene-level associations with multivariate GWAS of 35 mental traits. All significant gene sets are presented in ***supplementary table 8***. All p-values are corrected for multiple comparisons using Bonferroni correction.

### Boosting discovery of genetic loci associated with Big 5 personality traits and cognitive function

We used the conditional false discovery rate method (cFDR) (Smeland et al., 2019a) to leverage the additional power generated by our multivariate analysis to boost discovery of novel genetic loci associated with the remaining big 5 personality traits: agreeableness, conscientiousness, extraversion and openness in an independent sample (n=59,225) (Lo et al., 2017). cFDR applies a Bayesian model-free statistical framework to re-rank SNP associations with a primary trait given their strength of association with a conditional trait.

We identified novel loci associated agreeableness (n=11), conscientiousness (n=36), extraversion (n=89), and openness (n=24) (***figure 5a, supplementary tables 9-12***). This included, to our knowledge, the first genetic loci associated with agreeableness. The conditional analysis ensures that the boost in power from MOSTest method is driven by overlapping genetic variants, and not non-specific effects. Functional annotation of cFDR results identified 47 positionally-mapped genes for agreeableness, 157 for conscientiousness, 531 for extraversion, and 114 for openness (***supplementary tables 13-16***). Since MAGMA cannot be applied to cFDR statistics, we applied a hypergeometric test-based gene-set and tissue enrichment analyses using positionally mapped genes to replicate the approach taken by MAGMA (Watanabe et al., 2017). There were no gene-sets or tissues significantly enriched with mapped genes from any of the 4 traits.

**Figure 5:**
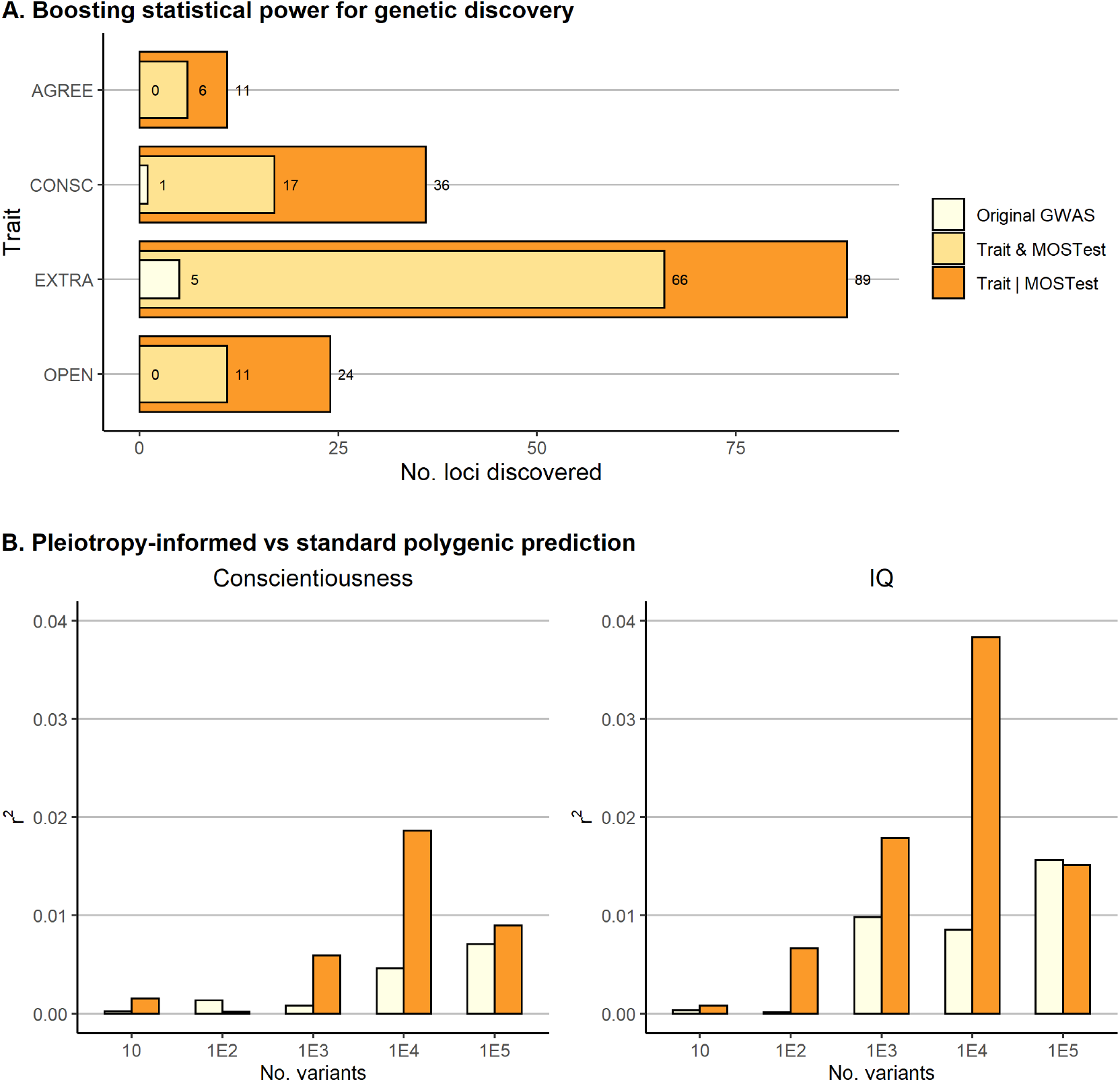
Leveraging multivariate analysis to boost discovery and polygenic prediction of personality and cognitive function. **A**. The number of loci associated with agreeableness (AGREE), conscientiousness (CONSC), extraversion (EXTRA), and openness (OPEN) in the primary GWAS (pale orange) compared to the conditional false discovery rate conditioning on the multivariate analysis of 35 mental measures (cFDR, dark orange). We also provide the number of shared genetic loci between personality and the multivariate analysis of 35 mental measures (conjFDR, orange). The number of loci discovered increased substantially, including the first loci reported for AGREE. **B**. Explained variance of CONSC and IQ polygenic scores (PGS) (r2, y-axis) for top 10, 100, 1000, 10,000 and 100,000 independent variants using primary GWAS p-value based ranking (light orange) and pleiotropy informed (cFDR-based) ranking (dark orange) from 23andMe and CHARGE GWAS summary statistics respectively. PGSs were tested in healthy participants from the Thematically Organised Psychosis study.

To test for pleiotropic effects in the remaining personality traits, we also performed conjunctional FDR (conjFDR), an extension of cFDR which identifies shared loci between two phenotypes. This revealed that 46-74% of loci associated with the Big 5 personality traits were also associated with our multivariate analysis of mental traits, indicating extensive pleiotropic effects beyond just neuroticism (***supplementary tables 17-20***).

We performed cFDR using independent neuroticism and general cognitive function GWAS, and compared these findings to the larger UKB-based GWAS to test the validity of cFDR in this context (***supplementary tables 21-22***). Of those present in both datasets, 50 out of 72 (69.4%) neuroticism and 92 out of 131 (70.2%) general cognitive function lead variants were nominally significant in the larger GWAS of neuroticism (Nagel et al., 2018a) and general intelligence (Savage et al., 2018), respectively.

### Improving polygenic prediction of personality and cognitive function

We investigated whether our multivariate GWAS could also improve polygenic prediction of Big 5 personality traits and cognitive function using a pleiotropy-informed PGS (pleioPGS) (van der Meer et al., 2020b). We constructed PGS using the 23andMe and CHARGE datasets for personality traits and general cognitive function, respectively, and tested their performance in Big 5 personality and IQ scores from healthy controls from the independent Thematically Organised Psychosis (TOP) study (n=578-1066). We compared the top 10-100,000 SNPs using original GWAS p-value ranking and cFDR-based ranking, hypothesising that the boost in power from our multivariate analysis will select more informative variants than standard GWAS, resulting in improved PGS performance. cFDR-based ranking outperformed original PGSs by 2.6 and 2.5 times for conscientiousness and IQ, respectively (***figure 5b***). PGSs performed poorly in the remaining four phenotypes for both p-value-based and cFDR-based ranking, indicating a lack of signal (max. r2 <0.01) (***supplementary figure 7***).

## Discussion

In this multivariate genome-wide association analysis of 35 heritable mental traits, we provide evidence of abundant pleiotropic genetic associations across personality and cognitive traits. Despite weak genetic and phenotypic correlations between neuroticism and cognitive domains, we discovered 431 genetic loci associated with the multivariate distribution of included traits, with evidence of pleiotropic associations across domains. Furthermore, we identified distinct patterns of relationships with evidence of cross-domain genetic association and mixed effect directions. Nonetheless, most lead SNPs were not genome-wide significant in univariate GWAS, demonstrating the boost in power provided by our multivariate approach. Functional characterisation revealed that the genetic signal captured by MOSTest was associated with increased gene expression across all brain tissues, the testis and ovary, and implicated synaptic structure and neurodevelopmental processes. We subsequently leveraged the extra power generated by our multivariate approach to boost discovery of genetic loci associated with the remaining Big 5 personality traits, identifying 160 loci for agreeableness (n=11), conscientiousness (n=36), extraversion (n=89), and openness (n=24). We further showed how the genetic loci shared across cognition and multiple personality traits improved polygenic prediction of conscientiousness and IQ in an independent sample. These findings have implications for how we conceptualise the neurobiology of personality and cognition, indicating that their genetic foundations are tightly interrelated. Dimensional, multivariate approaches which account for the complex set of interactions across domains are therefore better suited to fully elucidate the molecular mechanisms contributing to these fundamentally human traits.

Firstly, the boost in power generated by our combined analysis of neuroticism and cognitive measures, alongside our findings of shared genetic associations across domains, is consistent with the hypothesis that these two mental constructs are influenced by pleiotropic genetic variants. This builds on recent evidence that differences in brain structure and function are associated with a similar pattern of pleiotropic genetic effects (van der Meer et al., 2020a; Roelfs et al., 2022; Shadrin et al., 2021). As larger numbers of genetic loci associated with complex mental traits are discovered (Gandal and Geschwind, 2021), it is becoming increasingly apparent that individual genetic variants impact multiple, diverse traits, with few phenotype-specific variants. This represents a key conceptual advance which has several implications. Firstly, while large univariate GWAS have provided insights into the neurobiology of specific traits (Nagel et al., 2018a; Savage et al., 2018), future studies need to be aware of the lack of specificity of most variants associated with complex mental phenotypes. To fully characterise a given genetic variant, its effect should be evaluated beyond the specific phenotype of interest as it is likely to have pleiotropic effects across diverse domains (Karlsson Linnér et al., 2021; van der Meer et al., 2020a). Secondly, as statistical power increases, the relative effect size of a variant will likely be more informative with regards to specificity and relevance for a given phenotype than the presence or absence of a statistical association. In this respect, conventional GWAS may become less a tool for discovery and more focused on the precision of effect size estimates. Thirdly, as we have shown here, pleiotropic genetic effects can be leveraged to help boost the power for genetic discovery and polygenic prediction in related traits.

When comparing effect sizes of MOSTest discovered lead variants across included measures, there was also evidence of mixed effect directions between neuroticism and cognitive domains. This is consistent with the finding of minimal genetic correlation yet pleiotropic effects between these two domains. Genetic correlation is a genome-wide summary measure of the correlation of effect sizes between two phenotypes (Bulik-Sullivan et al., 2015a). It is therefore possible for two phenotypes to share large numbers of genetic variants but possess minimal correlation if there is a balance of shared variants with the same and opposite effect directions on the two phenotypes (Bahrami et al., 2021; Smeland et al., 2019b, 2020). Shared genetic variants with mixed effects reflect phenotypic findings that neuroticism does not significantly predict high school educational performance (Mammadov, 2021) or cognitive function in older adults (Wettstein et al., 2017). Nonetheless, “executive function” and “reaction time” clusters shared variants with the “worry” cluster and “fluid intelligence/memory” shared variants with the “depressed affect” cluster which were strongly discordant, despite weak negative genetic correlations. This suggests that MOSTest may prioritise variants which have more strongly aligned effect alleles in relation to the genome-wide average. Further, the recent findings of pleiotropic genetic effects on brain structure and function (van der Meer et al., 2021; Roelfs et al., 2022; Shadrin et al., 2021), as well as patterns of widespread gene expression across different brain regions (Hawrylycz et al., 2015) underscore the highly inter-related functions of brain regions and structures. Taken with our findings, this indicates that a complex interplay between heritable brain functions result in patterns of heritable, inter-related, higher-order mental traits which contribute to the core characteristics of an individual.

We used MAGMA to provide biological insights into the statistical associations captured by MOSTest. Firstly, tissue enrichment analysis showed significant enrichment in all included brain tissues (GTEx Consortium et al., 2017), underscoring the distributed nature of the genetic variants discovered. There were also several relevant gene-sets identified, including “observational learning”, “behavior” and “cognition”, alongside several gene-sets related to synaptic structure and function. Since MAGMA tests for enrichment of positionally mapped genes and so is not biased by the selection of tissue-specific eQTL databases, this indicates that MOSTest is capturing biologically plausible genes and is not driven by non-specific genetic overlap, helping to validate our findings. Furthermore, the diverse set of brain tissues identified, including cortical structures, sub-cortical structures, the midbrain and the hindbrain, support the broader concept of pleiotropic effects across the brain both on a structural and functional level (van der Meer et al., 2020c; Roelfs et al., 2022; Shadrin et al., 2021). It is also interesting to note that both the testis and ovary were significantly enriched, although to a lesser degree than brain tissues. Sex hormones can act in the brain to regulate gene transcription and interact directly with neurotransmitter systems (Hornung et al., 2020). They are also known to impact cognition, particularly verbal and visuospatial abilities (Sacher et al., 2013), and emotional regulation (Sundström-Poromaa, 2018), a core feature of neuroticism (Widiger and Oltmanns, 2017). Despite this, gonadal tissue was not significantly enriched in either the aforementioned general intelligence (Savage et al., 2018) or neuroticism GWAS (Nagel et al., 2018a). This may be the result of the additional power achieved using MOSTest.

Finally, we leveraged the boost in power from our multivariate analysis to improve discovery of genetic loci associated with agreeableness, conscientiousness, extraversion, and openness. This included, to the best of our knowledge, the first genetic loci reported for agreeableness. Genetic overlap between schizophrenia and neuroticism and openness has previously been reported using cFDR (Smeland et al., 2017). Interestingly, five of the six loci shared between schizophrenia and openness were also identified in our openness cFDR analysis. Nonetheless, larger samples are required to validate these findings. By re-ranking genetic variants according to the MOSTest-informed cFDR values, we also improved polygenic prediction of conscientiousness and IQ. As has previously been shown for schizophrenia and bipolar disorder (van der Meer et al., 2020b), the PGSs outperformed standard GWAS-based ranking despite using the same weightings, suggesting that this method prioritises more predictive variants. This approach is similar to other recent examples using multivariate to enhance discovery (Roelfs et al., 2022) and prediction (Baselmans et al., 2019; Ip et al., 2021). Nonetheless, PGSs for agreeableness, extraversion, neuroticism, and openness failed to achieve adequate prediction in our independent test sample. This may have been due to a lack of statistical power, the use of different personality scales for the training (John et al., 1991) and test samples (Costa and McCrae, 2008), or cultural differences between the American 23andMe sample (Lo et al., 2017) and the Norwegian, research-focussed TOP sample (Simonsen et al., 2011).

Among multivariate approaches, MOSTest was particularly well suited for the analysis of multiple personality and cognitive traits (van der Meer et al., 2020a). MOSTest is more flexible than CCA since it can handle differences in sample size across included phenotypes and is more computationally efficient for high dimensional data (van der Meer et al., 2020a). By using permuted individual-level genotypes, MOSTest also robustly controls for type 1 error. It is also important to note that MOSTest differs fundamentally from genomic structural equation modelling (SEM) (Grotzinger et al., 2019), another widely used multivariate GWAS method. While genomic SEM models the latent factor underlying a matrix of genetic correlations, MOSTest models the multivariate distribution of included variables. This means MOSTest can identify variants which are shared across phenotypes even if they have mixed effect directions on each trait, which has been shown for many brain-related mental traits (Bahrami et al., 2021; Hindley et al., 2021; O’Connell et al., 2021).

There were limitations to this study. Firstly, this analysis only included European-ancestry participants due to differences in linkage disequilibrium between ancestral groups and a lack of large, deeply phenotyped non-White European samples. Larger samples and new methods for trans-ancestral analysis are required to ensure the generalisability of these findings. Secondly, there were differences in sample size between measures. This means that the genetic associations captured by MOSTest are likely to be driven to a greater extent by measures with larger sample sizes and that z-score estimates for measures with smaller sample sizes may be less precise. Despite this, we showed statistically significant associations with measures from both domains, supporting our main finding of pleiotropic effects. Thirdly, we combined cognitive measures taken at different timepoints during the study. While systematic differences in cognitive performance may subtly alter the results, it is unlikely to change the main findings of the study. Fourthly, MOSTest requires the use of individual level data. This limited our ability to include other personality traits in the main analysis which were not included in UKB. We mitigated this by using our multivariate analysis to boost discovery for the remaining four personality traits. Finally, we used MAGMA for gene-mapping, tissue enrichment and gene-set analyses, which does not incorporate eQTL or chromatin interaction gene-mapping. This increased the specificity of the gene-mapping approach and meant that the gene-set and tissue enrichment analyses were not biased by the selection of eQTL or chromatin interaction databases. However, this also reduced the sensitivity of our gene-mapping procedure. We considered this approach to be the most appropriate since gene discovery was not an explicit aim of the present study.

## Conclusions

By combining 35 item and task-level measures of mental functioning in a multivariate framework, we demonstrate that distinct cognitive and personality traits are influenced by hundreds of genetic variants with pleiotropic effects and mixed effect directions, despite minimal genetic and phenotypic correlations. This contributes to a growing body of evidence indicating that common genetic variants underlying complex mental traits are closely interrelated, suggesting that “the whole is more than the sum of its parts” for brain-related phenotypes.

## Acknowledgements

We thank the research participants, employees and researchers of the UK Biobank, 23andMe, CHARGE and TOP for making this research possible. This work was partly performed on the TSD (Services for Sensitive Data) facilities, owned by the University of Oslo, operated and developed by the TSD service group at the University of Oslo, IT-Department (USIT). Computations were also performed on resources provided by UNINETT Sigma2—the National Infrastructure for High Performance Computing and Data Storage in Norway. We gratefully acknowledge support from the American National Institutes of Health (NS057198, EB00790), the Research Council of Norway (RCN) (229129, 213837, 324252, 300309, 273291, 223273), the South-East Norway Regional Health Authority (2022-073), KG Jebsen Stiftelsen (SKGJ-MED-021) and the US Norway Collaboration (RCN# 248980, 296030). This project has received funding from the European Union’s Horizon 2020 research and innovation programme under grant agreement No 847776 and 964874. The background vectors used in figure 1 were created by freepik - www.freepik.com and smart.servier.com.

## Author Contributions

Conceptualization, G.H., A.S., D.v.d.M., A.M.D. and O.A.A.; Methodology, AS, D.v.d.M, O.F., A.M.D.; Formal Analysis, A.S., G.H., N.P., W.C.; Resources, O.F., O.A.A.; Data Curation, A.S., D.v.d.M., O.F., O.B.S., Writing – Original Draft, G.H., A.S.; Writing – Review & Editing, All co-authors; Visualization, G.H., A.S.; Supervision, A.M.D., O.A.A.; Project Administration, O.A.A.; Funding Acquisition, A.M.D., O.A.A.

## STAR Methods

### Samples and phenotyping

#### UK Biobank

Genotypes, demographic, and clinical data were obtained from the UK Biobank. We selected unrelated (included in UKB genetic principal components calculation), white British individuals (as derived from both self-declared ethnicity and principal component analysis) with no sex chromosome aneuploidies (Bycroft et al., 2018) and genotyping call rate greater than 0.9. Participants who had withdrawn their consent were removed. This resulted in 337,145 individuals with mean age of 56.9 (standard deviation = 8.0 years). 53.7% were female. For the association analysis we retained only variants on autosomes with minor allele frequency above 0.001 imputation info score > 0.8 and with Hardy-Weinberg Equilibrium p-value > 1E-10, leaving 12.9 million variants.

***Table 1*** and ***supplementary table 1*** summarise the phenotypes included in our multivariate analysis. The UKB neuroticism items were derived from the Eysenck Personality Questionnaire-Revised Short Form (Eysenck et al., 1985). The scale was completed by all participants during enrolment at the assessment centre as part of the touchscreen assessment. All items comprised binary yes/no response options. Cognitive measures were collected at three different timepoints – either as part of the touchscreen cognitive assessment at enrolment (2006-2010), online cognitive follow-up (2014-2015) or during a follow-up imaging visit (2016). Some items from the touchscreen assessment were repeated during the cognitive follow-up and so were merged to maximise sample size. Included measures spanned a variety of cognitive domains, including verbal/numeric reasoning, prospective memory, working memory, non-verbal reasoning, visual declarative memory, processing speed and executive function (Fawns-Ritchie and Deary, 2020). All cognitive measures were coded so that larger values indicate better performance (i.e. shorter reaction time, less matching errors, faster task completion etc.).

#### 23andMe and CHARGE

For our replication and cFDR analyses, summary statistics for 23andMe Big 5 personality traits (Lo et al., 2017) and CHARGE general cognitive function (Davies et al., 2015) were accessed through collaborations. Sample make-up, genotyping procedures and phenotyping have been described in detail in the original publications (Davies et al., 2015; Lo et al., 2017). Briefly, the 23andMe samples comprised 59,225 individuals of European ancestry. Sum-scores for agreeableness, conscientiousness, extraversion, neuroticism, and openness were derived from the Big Five Inventory – 44-item edition (John et al., 1991). 23andMe customers completed the questionnaire online. The CHARGE general cognitive function sample comprised a meta-analysis of 53,949 participants of European ancestry from 31 cohorts. Cognitive function was assessed using a wide variety of different cognitive tests for fluid cognitive function. Each cohort included a minimum of three different tasks, and the principal component of included tasks for each cohort was computed to represent the “general cognitive function” phenotype.

#### TOP Sample

The TOP sample comprised participants recruited as healthy controls for an observational study of severe mental illness. Participants were identified at random from the national population register. Inclusion criteria included the absence of current or previous psychiatric disorder as identified by the Primary Care Evaluation of Mental Disorders (Prime-MD) delivered by a trained research assistant (Spitzer et al., 1994). Exclusion criteria were substance use disorder, physical health condition, previous traumatic brain injury, neurological disorders, autism spectrum disorder, personal or family (1^st^ degree relative) history of severe psychiatric disorder, and age outside of the range 13-72. Big 5 personality traits were assessed using the revised Neuroticism-Extraversion-Openness Five Factor Inventory (NEO-FFI) (Costa and McCrae, 1989), Norwegian edition, a 60-item questionnaire comprising 5-point Likert scale responses. IQ was measured using the Wechsler Abbreviated Scale of Intelligence second addition (WASI-II) (Wechsler, 1999). Incomplete responses were dropped, leaving sample sizes of 587 for agreeableness, 600 for conscientiousness, 581 for extraversion, 598 for neuroticism, 578 for openness and 1066 for IQ.

### Ethical considerations

All participants provided informed consent. UKB participants who withdrew consent were excluded from the study. UK Biobank data was accessed under accession number 27412. The 23andMe sample participated under a protocol approved by the external AAHRPP-accredited IRB, Ethical & Independent Review Services (E&I Review). Participants were included in the analysis on the basis of consent status as checked at the time data analyses were initiated. The use of summary statistics for cFDR analysis was evaluated by The Norwegian Institutional Review Board: Regional Committees for Medical and Health Research Ethics (REC) South-East Norway and found that no additional ethical approval was required because no individual data were used. TOP received ethical approval from Norwegian REC (ref. 2009/2485), Data Inspectorate (ref. 03/02051), and The Norwegian Directorate of Health (ref. 05/5821).

### Data analysis

#### Pre-processing of UKB variables

Prior to the association testing each item was manually pre-processed. Missing values were dropped from the analysis. Several continuous items with skewed and highly sparse distribution of answers were binarized. All continuous items were transformed using rank-based inverse normal transformation. Further details are provided in ***supplementary table 1***.

#### LD score regression heritability, genetic correlation, phenotypic correlation and hierarchical clustering

Univariate h^2^_SNP_ and pairwise genetic correlations (r_g_) were estimated using LDSR (Bulik-Sullivan et al., 2015a, 2015b). Briefly, LDSR estimates univariate h^2^_SNP_ from GWAS summary statistics by modelling the relationship between variant-level effect size and extent of LD, building on the observation that the larger the region of LD the larger the effect size estimate. Genetic correlation is then computed as the co-variance of SNP effect size between two traits after controlling for LD. We performed hierarchical clustering on pair-wise genetic correlations using Agglomerative Clustering algorithm with distance function 1-|r_g_|, as implemented in sklearn Python package (Pedregosa et al., 2011). Phenotypic correlations were computed using Spearman rank correlation as implemented in the Python package SciPy (Virtanen et al., 2020).

#### MOSTest and min-P

Plink2 (Purcell and Chang) was applied to perform item-level genotype-phenotype association testing using linear regression for continuous items and logistic regression for binary items with sex age and first 10 genetic principal components as covariates. In total we performed GWAS of 13 neuroticism and 26 cognition measures. Corresponding summary statistics were processed with LD score regression (Bulik-Sullivan et al., 2015b) to estimate SNP-heritabilities (***supplementary table 2, supplementary figure 1***) and genetic correlations between items (***figure 1***). Since including non-heritable traits into MOSTest analysis may reduce statistical power (van der Meer et al., 2020a), only items with h^2^ p-value < 3.167E-5 were used for subsequent MOSTest and min-P analyses. This threshold recommended by the developers of LDSR-based SNP-heritability (Bulik-Sullivan et al., 2015b) and has previously been used for large-scale heritability analyses of UKB genetic data (Walters). In total 35 measures (13 neuroticism and 22 cognition cognitive) passed this h^2^_SNP_ filter. Variant z-scores from item and task-level GWAS for these 35 measures were combined in MOSTest and min-P analyses to produce multivariate p-values as described elsewhere (Shadrin et al., 2021). For MOSTest we selected regularization parameter (r=3) which provided the largest yield of genetic loci (Shadrin et al., 2021). We also performed MOSTest analyses for only neuroticism measures and only cognitive measures.

Genetic overlap between MOSTest across univariate GWAS analyses was determined at the lead-variant level. We extracted p-values for all MOSTest lead variants from each individual univariate GWAS for included measures. Genetic overlap was deemed present if the lead variant was significant in each pair of univariate GWAS at the specified significance threshold (p<5×10^−8^, p<1×10^−6^, p<1×10^− 5^). The same procedure was used to quantify overlap across the three multivariate analyses.

We performed hierarchical clustering of univariate z-scores for each MOSTest-discovered lead variant. Hierarchical clustering was produced using AgglomerativeClustering algorithm with Euclidian distance, as implemented in sklearn Python package. Lead variants were split into 7 clusters. For each variable we then estimated the median z-score over all variants in the cluster.

#### Conditional/conjunctional false discovery rate

We applied cFDR to boost discovery of genetic variants associated with the Big 5 personality traits and general cognitive function. Firstly, conditional qq-plots were constructed by comparing enrichment of association in all variants in the primary trait (i.e Big 5 personality traits or general cognitive function) with 3 subsets of variants defined by their strength of association (p<0.1, p<0.01 and p<0.001) with the secondary trait (i.e. MOSTest summary statistics). Successive left-ward deflection with increasing threshold of significance indicates cross-trait enrichment. Shift in enrichment conditional on the secondary trait can be directly interpreted according to the Bayesian definition of the true discovery rate (TDR = 1-FDR), whereby a larger shift is consistent with a smaller FDR. This means cFDR values can be computed for each variant by comparing enrichment of all variants with a subset of variants which are as strongly or more strongly associated with the secondary trait. The cFDR value can therefore be interpreted as the probability that a given SNP is not associated with the primary trait given that the SNP is more strongly or as strongly associated with both phenotypes than observed in the original GWAS. Look-up plots can therefore be constructed which provide cFDR values given the p-values in the primary and secondary traits. Conjunctional FDR statistic is subsequently computed by repeating the analysis having switched the primary and secondary trait. The maximum of the two cFDR statistics represents the probability that a given SNP is not associated with the primary or secondary trait given that the SNP is more strongly or as strongly associated with both phenotypes than observed in the original GWAS. We performed 100 iterations of each analysis after random pruning from independent LD blocks (r^2^>0.1). Genomic inflation was corrected for by a conservative genomic control procedure utilizing intergenic variants which lack true associations relative to other functional regions (Schork et al., 2013). The MHC region was excluded from the model-fitting procedure to prevent inflation of test statistics due to complex LD.

### Locus definition

Genetic loci were defined based on association summary statistics produced with MOSTest, min-P and cFDR following the protocol implemented in FUMA with default parameters (Watanabe et al., 2017). The protocol is summarised as follows:

1. Independent significant genetic variants were identified as variants with p-value<5E-8 or cFDR<0.05 and linkage disequilibrium (LD) r2<0.6 with each other.
2. A subset of these independent significant variants with LD r2<0.1 were selected as lead variants.
3. For each independent significant variant all candidate variants were identified as variants with LD r2≥0.6.
4. For a given lead variant the borders of the genomic locus were defined as min/max positional coordinates over all corresponding candidate variants.
5. Loci were merged if they were separated by less than 250kb.

### Replication in independent samples

We tested for *en masse* sign concordance of genetic effects in MOSTest-discovered lead SNPs between UKB fluid intelligence sum-score and CHARGE general cognitive function summary statistics, and UKB neuroticism sum-score and 23andMe neuroticism summary statistics. We dropped all variants which were not present in the independent summary statistics and variants with ambivalent effect alleles. We first used an exact binomial test to test the null hypothesis that sign concordance was randomly distributed (p=0.5), given the total number of variants (n) and the number of variants with concordant effects in UKB and each independent dataset, respectively (k). To test for evidence of pleiotropic effects, we used an exact binomial test to test the null hypothesis that sign concordance in both neuroticism and cognitive function were randomly distributed (p=0.25), given the total number of variants (n) and the number of variants which were concordant in both phenotypes simultaneously. We also extracted p-values from the primary GWASs and reported the number of nominally significant variants in independent samples.

### Mapped genes, tissue specificity and gene-set analyses

Gene-mapping of MOSTest GWAS summary statistics were performed using MAGMA as implemented in FUMA. The MHC region was excluded and all other settings were default. Gene analyses of individual items (***supplementary* results**) were performed with MAGMA (version 1.09b) (de Leeuw et al., 2015) applying a SNP-wide mean model to GWAS summary statistics excluding variants within MHC region (chr6:25000000-33000000) 1000 Genomes Phase 3 EUR used as a reference panel and other settings being default. 18,952 genes were included in the analysis.

Tissue specificity and gene-set enrichment analysis of MOSTest summary statistics was performed using MAGMA as implemented in FUMA (de Leeuw et al., 2015). Tissue specificity was tested in GTEx version 7 eQTL database (GTEx Consortium et al., 2017) across 53 “detail tissues” and 30 “general tissues”. Gene-set enrichment was tested in Gene Ontology (Ashburner et al., 2000) and curated gene-sets from MsigDB (Liberzon et al., 2011) (n=10,678). Bonferroni correction was applied to correct for multiple comparisons.

Since cFDR statistics are not applicable to MAGMA, genes were mapped to candidate SNPs by positional mapping, i.e. according to their physical proximity (<10kb) to each variant. We performed tissue specificity and gene-set analysis using the GENE2FUNC functionality in FUMA using default settings. Positionally mapped genes were used as input for all analyses. Over-representation of mapped genes within tissue-specific differentially expressed genes, and Gene Ontology and curated gene-sets was tested using a hypergeometric test. Correction for multiple comparisons was performed using the Benjamini-Hochberg method.

### Polygenic score analysis

PGSs were constructed using PRSice (Choi and O’Reilly, 2019) from summary statistics from 23andMe Big 5 personality traits (Lo et al., 2017) and CHARGE general cognitive function (Davies et al., 2015). Using the pleioPGS approach (van der Meer et al., 2020b), we leveraged our multivariate analysis by comparing standard GWAS-ranked lead SNPs with cFDR-based ranking, using the same weights derived from the original GWAS (Baselmans et al., 2019; Ip et al., 2021; van der Meer et al., 2020b). Sex, age and 20 principal components were included as covariates. cFDR and PGS plots were generated using the ggplot2 package in r as implemented in rstudio (Allaire, 2012; Team, 2013; Wickham, 2016).

## Data availability

Individual-level UKB data is available through a publicly accessible application via UKB (https://www.ukbiobank.ac.uk/enable-your-research/apply-for-access). The full GWAS summary statistics for the 23andMe discovery data set will be made available through 23andMe to qualified researchers under an agreement with 23andMe that protects the privacy of the 23andMe participants. Please visit https://research.23andme.com/collaborate/=dataset-access/ for more information and to apply to access the data. CHARGE general cognitive function summary statistics are publicly available at https://www.chargeconsortium.com/main/results.

## Code availability

Code for MOSTest and cFDR are publicly available at https://github.com/precimed/mostest/tree/mental and https://github.com/precimed/pleiofdr.

## Supplementary Information

1. Supplementary material: supplementary results and supplementary figures 1-9
2. Supplementary tables: Supplementary tables 1-22

## References

Allaire, J. (2012). RStudio: integrated development environment for R. Boston, MA 770, 165–171.

Andreassen, O.A., Thompson, W.K., Schork, A.J., Ripke, S., Mattingsdal, M., Kelsoe, J.R., Kendler, K.S., O’Donovan, M.C., Rujescu, D., and Werge, T. (2013). Improved detection of common variants associated with schizophrenia and bipolar disorder using pleiotropy-informed conditional false discovery rate. PLoS Genetics 9, e1003455.

Ashburner, M., Ball, C.A., Blake, J.A., Botstein, D., Butler, H., Cherry, J.M., Davis, A.P., Dolinski, K., Dwight, S.S., Eppig, J.T., et al. (2000). Gene Ontology: tool for the unification of biology. Nature Genetics 25, 25–29.

Bahrami, S., Hindley, G., Winsvold, B.S., O’Connell, K.S., Frei, O., Shadrin, A., Cheng, W., Bettella, F., Rødevand, L., Odegaard, K.J., et al. (2021). Dissecting the shared genetic basis of migraine and mental disorders using novel statistical tools. Brain.

Baselmans, B.M.L., Jansen, R., Ip, H.F., van Dongen, J., Abdellaoui, A., van de Weijer, M.P., Bao, Y., Smart, M., Kumari, M., Willemsen, G., et al. (2019). Multivariate genome-wide analyses of the well-being spectrum. Nature Genetics 2019 51:3 51, 445–451.

Bulik-Sullivan, B., Finucane, H.K., Anttila, V., Gusev, A., Day, F.R., Loh, P.-R., Duncan, L., Perry, J.R.B., Patterson, N., and Robinson, E.B. (2015a). An atlas of genetic correlations across human diseases and traits. Nature Genetics 47, 1236.

Bulik-Sullivan, B.K., Loh, P.-R., Finucane, H.K., Ripke, S., Yang, J., Patterson, N., Daly, M.J., Price, A.L., Neale, B.M., and Consortium, S.W.G. of the P.G. (2015b). LD Score regression distinguishes confounding from polygenicity in genome-wide association studies. Nature Genetics 47, 291.

Bycroft, C., Freeman, C., Petkova, D., Band, G., Elliott, L.T., Sharp, K., Motyer, A., Vukcevic, D., Delaneau, O., and O’Connell, J. (2018). The UK Biobank resource with deep phenotyping and genomic data. Nature 562, 203–209.

Choi, S.W., and O’Reilly, P.F. (2019). PRSice-2: Polygenic Risk Score software for biobank-scale data. GigaScience 8, giz082.

Costa, P.T., and McCrae, R.R. (1989). NEO five-factor inventory (NEO-FFI). Odessa, FL: Psychological Assessment Resources 3.

Costa, P.T., and McCrae, R.R. (2008). The revised NEO personality inventory (NEO-PI-R). The SAGE Handbook of Personality Theory and Assessment: Volume 2 - Personality Measurement and Testing 179–198.

Cullen, B., Smith, D.J., Deary, I.J., Evans, J.J., and Pell, J.P. (2017). The ‘cognitive footprint’ of psychiatric and neurological conditions: cross-sectional study in the UK Biobank cohort. Acta Psychiatrica Scandinavica 135, 593–605.

Damian, R.I., Spengler, M., Sutu, A., and Roberts, B.W. (2019). Sixteen going on sixty-six: A longitudinal study of personality stability and change across 50 years. Journal of Personality and Social Psychology 117, 674–695.

Davies, G., Armstrong, N., Bis, J.C., Bressler, J., Chouraki, V., Giddaluru, S., Hofer, E., Ibrahim-Verbaas, C.A., Kirin, M., Lahti, J., et al. (2015). Genetic contributions to variation in general cognitive function: a meta-analysis of genome-wide association studies in the CHARGE consortium (N=53949). Molecular Psychiatry 20, 183–192.

Eysenck, S.B.G., Eysenck, H.J., and Barrett, P. (1985). A revised version of the psychoticism scale. Personality and Individual Differences 6, 21–29.

Fawns-Ritchie, C., and Deary, I.J. (2020). Reliability and validity of the UK Biobank cognitive tests. PloS One 15.

Gandal, M.J., and Geschwind, D.H. (2021). Polygenicity in Psychiatry—Like It or Not, We Have to Understand It. Biological Psychiatry 89, 2–4.

Grotzinger, A.D., Rhemtulla, M., de Vlaming, R., Ritchie, S.J., Mallard, T.T., Hill, W.D., Ip, H.F., Marioni, R.E., McIntosh, A.M., Deary, I.J., et al. (2019). Genomic SEM Provides Insights into the Multivariate Genetic Architecture of Complex Traits. Nature Human Behaviour 3, 513.

GTEx, Consortium, Aguet, F., Brown, A.A., Castel, S.E., Davis, J.R., He, Y., Jo, B., Mohammadi, P., Park, Y., Parsana, P., et al. (2017). Genetic effects on gene expression across human tissues. Nature 550, 204.

Hawrylycz, M., Miller, J.A., Menon, V., Feng, D., Dolbeare, T., Guillozet-Bongaarts, A.L., Jegga, A.G., Aronow, B.J., Lee, C.K., Bernard, A., et al. (2015). Canonical genetic signatures of the adult human brain. Nature Neuroscience 18, 1832–1844.

Hill, W.D., Weiss, A., Liewald, D.C., Davies, G., Porteous, D.J., Hayward, C., McIntosh, A.M., Gale, C.R., and Deary, I.J. (2020). Genetic contributions to two special factors of neuroticism are associated with affluence, higher intelligence, better health, and longer life. Molecular Psychiatry 25, 3034–3052.

Hindley, G., Bahrami, S., Steen, N.E., O’Connell, K.S., Frei, O., Shadrin, A., Bettella, F., Rødevand, L., Fan, C.C., Dale, A.M., et al. (2021). Characterising the shared genetic determinants of bipolar disorder, schizophrenia and risk-taking. Translational Psychiatry 11.

Hornung, J., Lewis, C.A., and Derntl, B. (2020). Sex hormones and human brain function. Handbook of Clinical Neurology 175, 195–207.

Ip, H.F., van der Laan, C.M., Krapohl, E.M.L., Brikell, I., Sánchez-Mora, C., Nolte, I.M., St Pourcain, B., Bolhuis, K., Palviainen, T., Zafarmand, H., et al. (2021). Genetic association study of childhood aggression across raters, instruments, and age. Translational Psychiatry 2021 11:1 11, 1–9.

John, O.P., Donahue, E.M., and Kentle, R. (1991). The big five inventory: Versions 4a and 54 [Technical Report]. Berkeley: University of California, Institute of Personality and Social Research.

Karlsson Linnér, R., Mallard, T.T., Barr, P.B., Sanchez-Roige, S., Madole, J.W., Driver, M.N., Poore, H.E., de Vlaming, R., Grotzinger, A.D., Tielbeek, J.J., et al. (2021). Multivariate analysis of 1.5 million people identifies genetic associations with traits related to self-regulation and addiction. Nature Neuroscience 24, 1367–1376.

Lee, J.J., Wedow, R., Okbay, A., Kong, E., Maghzian, O., Zacher, M., Nguyen-Viet, T.A., Bowers, P., Sidorenko, J., Karlsson Linnér, R., et al. (2018). Gene discovery and polygenic prediction from a genome-wide association study of educational attainment in 1.1 million individuals. Nature Genetics 50, 1112–1121.

de, Leeuw, C.A., Mooij, J.M., Heskes, T., and Posthuma, D. (2015). MAGMA: Generalized Gene-Set Analysis of GWAS Data. PLoS Computational Biology 11.

Liberzon, A., Subramanian, A., Pinchback, R., Thorvaldsdóttir, H., Tamayo, P., and Mesirov, J.P. (2011). Molecular signatures database (MSigDB) 3.0. Bioinformatics 27, 1739–1740.

Lo, M.-T., Hinds, D.A., Tung, J.Y., Franz, C., Fan, C.-C., Wang, Y., Smeland, O.B., Schork, A., Holland, D., Kauppi, K., et al. (2017). Genome-wide analyses for personality traits identify six genomic loci and show correlations with psychiatric disorders. Nature Genetics 49, 152–156.

Mammadov, S. (2021). Big Five personality traits and academic performance: A meta-analysis. Journal of Personality.

van der Meer, D., Frei, O., Kaufmann, T., Shadrin, A.A., Devor, A., Smeland, O.B., Thompson, W.K., Fan, C.C., Holland, D., Westlye, L.T., et al. (2020a). Understanding the genetic determinants of the brain with MOSTest. Nature Communications 11, 3512.

van der Meer, D., Shadrin, A.A., O’connell, K., Bettella, F., Djurovic, S., Wolfers, T., Alnaes, D., Agartz, I., Smeland, O.B., Melle, I., et al. (2020b). Improved prediction of schizophrenia by leveraging genetic overlap with brain morphology. MedRxiv 2020.08.03.20167510.

van der Meer, D., Rokicki, J., Kaufmann, T., Córdova-Palomera, A., Moberget, T., Alnæs, D., Bettella, F., Frei, O., Doan, N.T., Sønderby, I.E., et al. (2020c). Brain scans from 21,297 individuals reveal the genetic architecture of hippocampal subfield volumes. Molecular Psychiatry 25, 3053–3065.

van der Meer, D., Kaufmann, T., Shadrin, A.A., Makowski, C., Frei, O., Roelfs, D., Monereo-Sánchez, J., Linden, D.E.J., Rokicki, J., Alnæs, D., et al. (2021). The genetic architecture of human cortical folding. Science Advances 7.

Nagel, M., Jansen, P.R., Stringer, S., Watanabe, K., de Leeuw, C.A., Bryois, J., Savage, J.E., Hammerschlag, A.R., Skene, N.G., Muñoz-Manchado, A.B., et al. (2018a). Meta-analysis of genome-wide association studies for neuroticism in 449,484 individuals identifies novel genetic loci and pathways. Nature Genetics 50, 920–927.

Nagel, M., Watanabe, K., Stringer, S., Posthuma, D., and van der Sluis, S. (2018b). Item-level analyses reveal genetic heterogeneity in neuroticism. Nature Communications 9.

O’Connell, K.S., Frei, O., Bahrami, S., Smeland, O.B., Bettella, F., Cheng, W., Chu, Y., Hindley, G., Lin, A., and Shadrin, A. (2021). Characterizing the genetic overlap between psychiatric disorders and sleep-related phenotypes. Biological Psychiatry 90, 621–631.

Pedregosa, F., Varoquaux, G., Gramfort, A., Michel, V., Thirion, B., Grisel, O., Blondel, M., Prettenhofer, P., Weiss, R., and Dubourg, V. (2011). Scikit-learn: Machine learning in Python. The Journal of Machine Learning Research 12, 2825–2830.

Purcell, S., and Chang, C. PLINK2 (v1. 90b6. 9). Available Online: Www.Cog-Genomics. Org/Plink/2.0/(Accessed on 1 May 2019).

Ripke, S., Walters, J.T.R., and Donovan, M.C. (2020). Mapping genomic loci prioritises genes and implicates synaptic biology in schizophrenia. MedRxiv 2020.09.12.20192922.

Roelfs, D., Meer, D. van der, Alnæs, D., Frei, O., Shadrin, A.A., Loughnan, R., Fan, C.C., Dale, A.M., Andreassen, O.A., Westlye, L.T., et al. (2022). Genetic overlap between multivariate measures of human functional brain connectivity and psychiatric disorders. MedRxiv 2021.06.15.21258954.

Sacher, J., Okon-Singer, H., and Villringer, A. (2013). Evidence from neuroimaging for the role of the menstrual cycle in the interplay of emotion and cognition. Frontiers in Human Neuroscience 7, 374.

Savage, J.E., Jansen, P.R., Stringer, S., Watanabe, K., Bryois, J., de Leeuw, C.A., Nagel, M., Awasthi, S., Barr, P.B., Coleman, J.R.I., et al. (2018). Genome-wide association meta-analysis in 269,867 individuals identifies new genetic and functional links to intelligence. Nature Genetics 50, 912–919.

Schork, A.J., Thompson, W.K., Pham, P., Torkamani, A., Roddey, J.C., Sullivan, P.F., Kelsoe, J.R., O’Donovan, M.C., Furberg, H., and Schork, N.J. (2013). All SNPs are not created equal: genome-wide association studies reveal a consistent pattern of enrichment among functionally annotated SNPs. PLoS Genet 9, e1003449.

Shadrin, A.A., Kaufmann, T., van der Meer, D., Palmer, C.E., Makowski, C., Loughnan, R., Jernigan, T.L., Seibert, T.M., Hagler, D.J., Smeland, O.B., et al. (2021). Vertex-wise multivariate genome-wide association study identifies 780 unique genetic loci associated with cortical morphology. NeuroImage 244, 118603.

Simonsen, C., Sundet, K., Vaskinn, A., Birkenaes, A.B., Engh, J.A., Faerden, A., Jónsdóttir, H., Ringen, P.A., Opjordsmoen, S., Melle, I., et al. (2011). Neurocognitive dysfunction in bipolar and schizophrenia spectrum disorders depends on history of psychosis rather than diagnostic group. Schizophrenia Bulletin 37, 73–83.

Smeland, O.B., Wang, Y., Lo, M.T., Li, W., Frei, O., Witoelar, A., Tesli, M., Hinds, D.A., Tung, J.Y., Djurovic, S., et al. (2017). Identification of genetic loci shared between schizophrenia and the Big Five personality traits. Scientific Reports 2017 7:1 7, 1–9.

Smeland, O.B., Frei, O., Shadrin, A., O’Connell, K., Fan, C.-C., Bahrami, S., Holland, D., Djurovic, S., Thompson, W.K., and Dale, A.M. (2019a). Discovery of shared genomic loci using the conditional false discovery rate approach. Human Genetics 1–10.

Smeland, O.B., Bahrami, S., Frei, O., Shadrin, A., O’Connell, K., Savage, J., Watanabe, K., Krull, F., Bettella, F., Steen, N.E., et al. (2019b). Genome-wide analysis reveals extensive genetic overlap between schizophrenia, bipolar disorder, and intelligence. Molecular Psychiatry 25, 844–853.

Smeland, O.B., Frei, O., Dale, A.M., and Andreassen, O.A. (2020). The polygenic architecture of schizophrenia — rethinking pathogenesis and nosology. Nature Reviews Neurology 16, 366–379.

Spitzer, R.L., Williams, J.B.W., Kroenke, K., Linzer, M., deGruy, F.V., Hahn, S.R., Brody, D., and Johnson, J.G. (1994). Utility of a new procedure for diagnosing mental disorders in primary care: the PRIME-MD 1000 study. Jama 272, 1749–1756.

Strickhouser, J.E., Zell, E., and Krizan, Z. (2017). Does personality predict health and well-being? A metasynthesis. Health Psychology 36, 797–810.

Sundström-Poromaa, I. (2018). The menstrual cycle influences emotion but has limited effect on cognitive function. Vitamins and Hormones 107, 349–376.

Team, R.C. (2013). R: A language and environment for statistical computing.

Virtanen, P., Gommers, R., Oliphant, T.E., Haberland, M., Reddy, T., Cournapeau, D., Burovski, E., Peterson, P., Weckesser, W., Bright, J., et al. (2020). SciPy 1.0: fundamental algorithms for scientific computing in Python. Nature Methods 17, 261–272.

Walhovd, K.B., Krogsrud, S.K., Amlien, I.K., Bartsch, H., Bjørnerud, A., Due-Tønnessen, P., Grydeland, H., Hagler, D.J., Håberg, A.K., Kremen, W.S., et al. (2016). Neurodevelopmental origins of lifespan changes in brain and cognition. Proceedings of the National Academy of Sciences 113, 9357–9362.

Walters, R. Heritability of >4,000 traits & disorders in UK Biobank.

Watanabe, K., Taskesen, E., van Bochoven, A., and Posthuma, D. (2017). Functional mapping and annotation of genetic associations with FUMA. Nature Communications 8, 1826.

Wechsler, D. (1999). Wechsler abbreviated scale of intelligence--.

Wettstein, M., Tauber, B., Kuźma, E., and Wahl, H.W. (2017). The interplay between personality and cognitive ability across 12 Years in middle and late adulthood: Evidence for reciprocal associations. Psychology and Aging 32, 259–277.

Wickham, H. (2016). ggplot2: elegant graphics for data analysis (springer).

Widiger, T.A., and Oltmanns, J.R. (2017). Neuroticism is a fundamental domain of personality with enormous public health implications. World Psychiatry 16, 144.

Wraw, C., Deary, I.J., Gale, C.R., and Der, G. (2015). Intelligence in youth and health at age 50. Intelligence 53, 23–32.

