## Supplementary material for "Multivariate genetic analysis of personality and cognitive traits reveals abundant pleiotropy and improves prediction"

Guy Hindley (M.D.)^1,2*†^, Alexey Shadrin (Ph.D.)^1,3*†^, Dennis van der Meer (Ph.D.)^1,4^, Nadine Parker (Ph.D.)^1^, Weiqiu Cheng (Ph.D.)^1^, Kevin S. O’Connell (Ph.D.)^1^, Shahram Bahrami (Ph.D.) ^1^, Aihua Lin (Ph.D.)^1^, Naz Karadag (M.Sc.)^1^, Børge Holen (M.D.)^1^, Chun C Fan (M.D.)^5,6^, Torill Ueland (Ph.D.) ^1,7^, Srdjan Djurovic (Ph.D.)^3,8,9^, Olav B. Smeland (M.D.)^1^, Oleksandr Frei (Ph.D.)^1,10^, Anders M. Dale (Ph.D.)^5,11,12,13^, Ole A. Andreassen (M.D.)^1,3*^

^1^NORMENT Centre, Institute of Clinical Medicine, University of Oslo and Division of Mental Health and Addiction, Oslo University Hospital, 0407 Oslo, Norway

^2^Psychosis Studies, Institute of Psychiatry, Psychology and Neurosciences, King’s College London, 16 De Crespigny Park, London SE5 8AB, United Kingdom

^3^KG Jebsen Centre for Neurodevelopmental disorders, University of Oslo, Oslo, Norway

^4^School of Mental Health and Neuroscience, Faculty of Health, Medicine and Life Sciences, Maastricht University, Maastricht, The Netherlands

^5^Department of Cognitive Science, University of California, San Diego, La Jolla, CA, USA

^6^Multimodal Imaging Laboratory, University of California San Diego, La Jolla, CA 92093, USA

^7^Department of Psychology, University of Oslo, Norway

^8^Department of Medical Genetics, Oslo University Hospital, Oslo, Norway

^9^NORMENT Centre, Department of Clinical Science, University of Bergen, Bergen, Norway

^10^Center for Bioinformatics, Department of Informatics, University of Oslo, PO box 1080, Blindern, 0316 Oslo, Norway

^11^Department of Psychiatry, University of California, San Diego, La Jolla, CA, USA

^12^Department of Neurosciences, University of California San Diego, La Jolla, CA 92093, USA

^13^Department of Radiology, University of California, San Diego, La Jolla, CA 92093, USA

*Corresponding authors

^†^These authors contributed equally

Correspondance:	Guy Hindley –

Alexey Shadrin –

Ole A Andreassen –

### Supplementary Results

#### Gene-level overlap despite minimal genetic and phenotypic correlations

We performed LDSR genetic correlation and Spearman rank phenotypic correlation on all pairs of heritable measures (***figure 1, supplementary table 2***). This revealed that neuroticism and cognitive measures were strongly positively genetically correlated within their respective domains (mean r_g_ = 0.65 and 0.56, respectively), with 100% (78/78) of neuroticism pairs and 93% (215/231) of cognitive pairs significantly correlated after Bonferroni correction. However, neuroticism items were weakly negatively correlated with cognitive measures (mean r_g_ = -0.14) and the majority were non-significant after Bonferroni correction (162/286).

There was moderate phenotypic correlation within neuroticism and cognitive measures (mean r_p_ = 0.34 and 0.17, respectively) and all pairwise analyses were significant after Bonferroni correction for both sets of pairwise analyses. In contrast, there were weaker negative correlations between neuroticism and cognitive measures (mean r_p_ = -0.03), and a larger proportion of non-significant correlations (67/286) (***supplementary table 2***).

#### Gene-level overlap between neuroticism and cognition supports evidence of mixed effect directions

The pattern of pleiotropic genetic effects was further evidenced by gene-level overlap between domains, despite the minimal genetic correlation. Based on the five genetic correlation-based clusters, we compared the number of overlapping MAGMA-mapped genes across each cluster. This revealed that the largest number of overlapping mapped genes was observed between neuroticism clusters (50.3%). However, the overlap across the neuroticism cluster “depressed affect” and cognitive cluster “executive function” (29.8%) was greater than the overlap within cognitive clusters (1.9%-27.7%) (***supplementary figure 3***). This pattern of gene-level overlap despite low genetic correlation is consistent with distributed genetic effects with mixed effect directions.

### Supplementary Figures

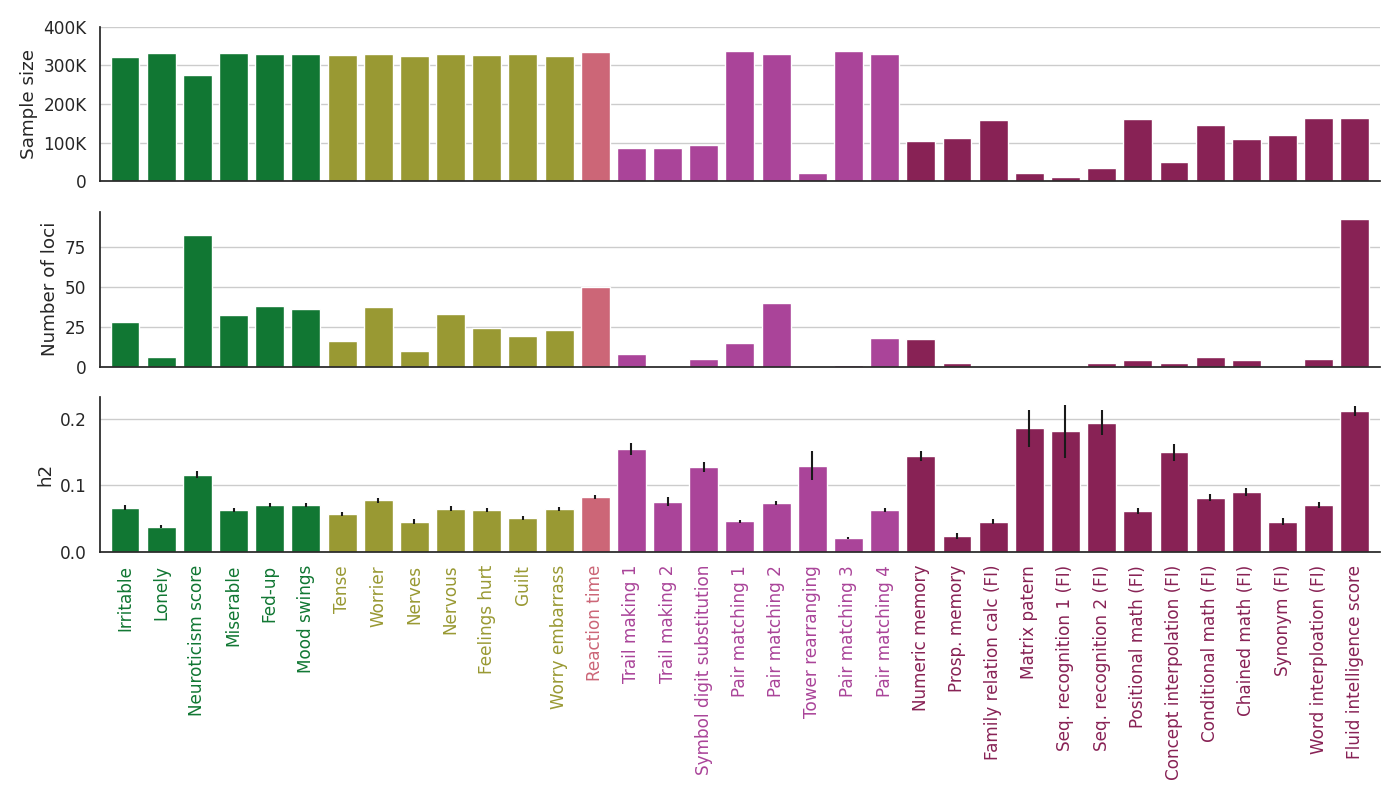

**Supplementary figure 1. Sample size, univariate GWAS discovery and SNP-heritability of neuroticism and cognitive measures**. Neuroticism measures (dark and light blue bars) had larger sample sizes than most cognitive measures (green, orange, and pink bars) which, in combination with SNP-heritability (h2), impacted the number of loci discovered for each measure. Please see **supplementary table 1** for a detailed description of each measure.

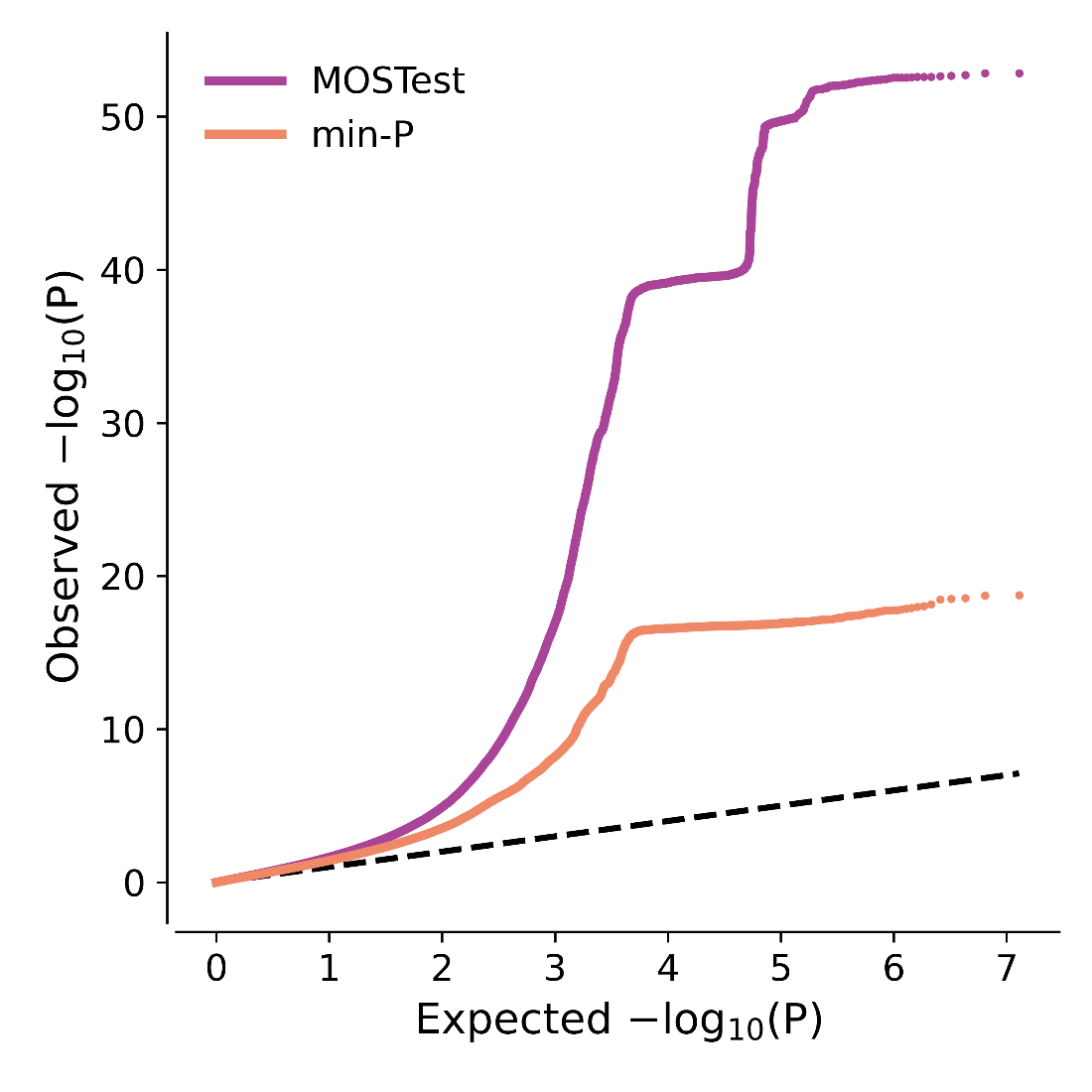

**Supplementary figure 2:** QQ plot comparing enrichment of MOSTest and min-P SNP associations. Observed -log_10_p-values (y axis) plotted against expected -log_10_p-values (x-axis). No statistical association is represented by the dashed line. The leftwards deflection of MOSTest SNP associations from the dashed line and the higher maximum observed -log_10_(p-value) indicates stronger enrichment and therefore a boost in power under the MOSTest statistical framework.

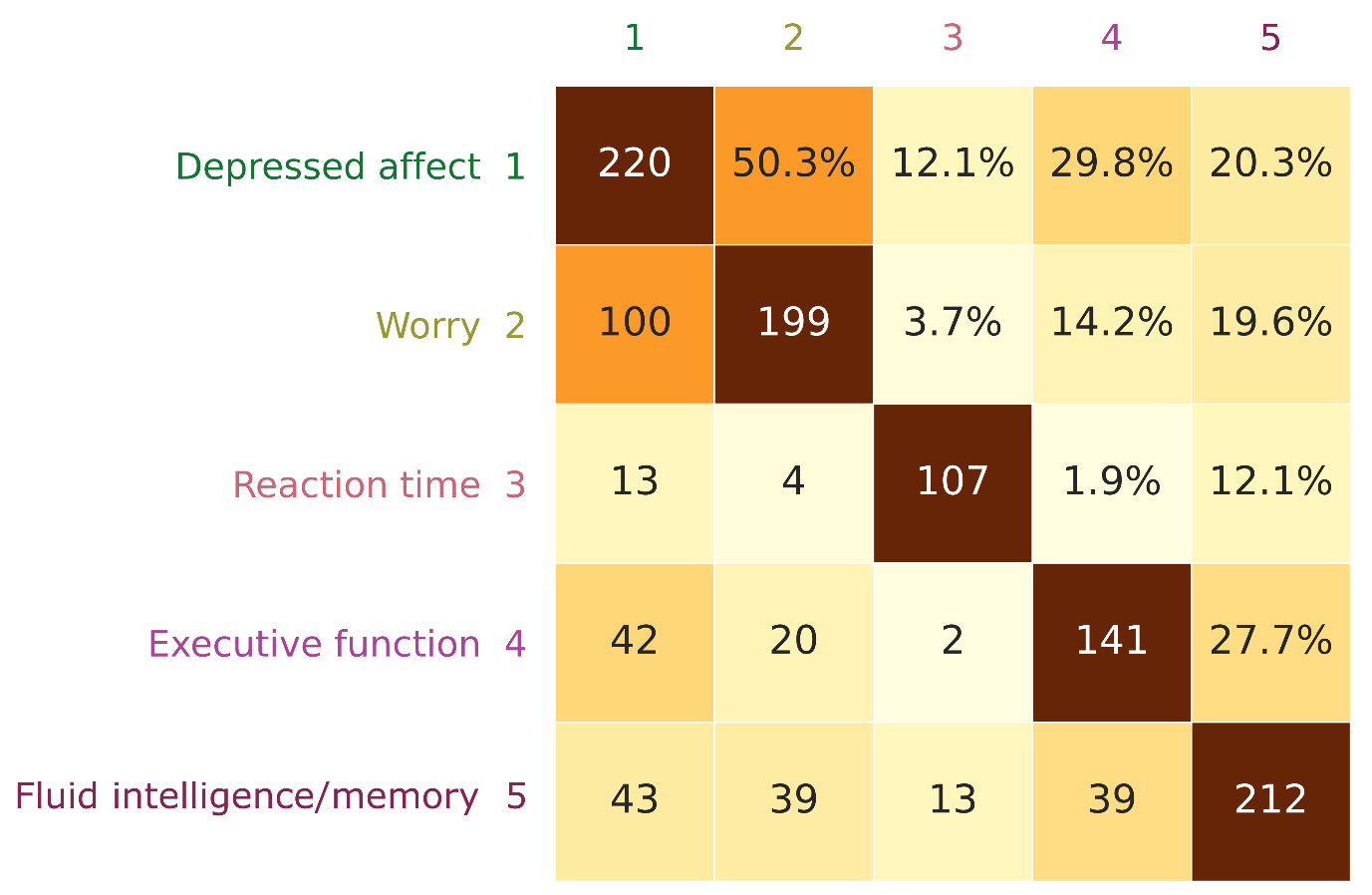

**Supplementary figure 3. Gene-level overlap across neuroticism and cognitive clusters.** Extensive overlap of MAGMA-mapped genes across neuroticism and cognitive clusters, both within and across domains. Diagonal: Total number of MAGMA-mapped genes for each cluster. Bottom left: The number of MAGMA-mapped genes from univariate GWAS overlapping across each pair of clusters. Top left: The proportion of MAGMA-mapped genes overlapping across each pair of clusters relative to the trait with fewer mapped genes.

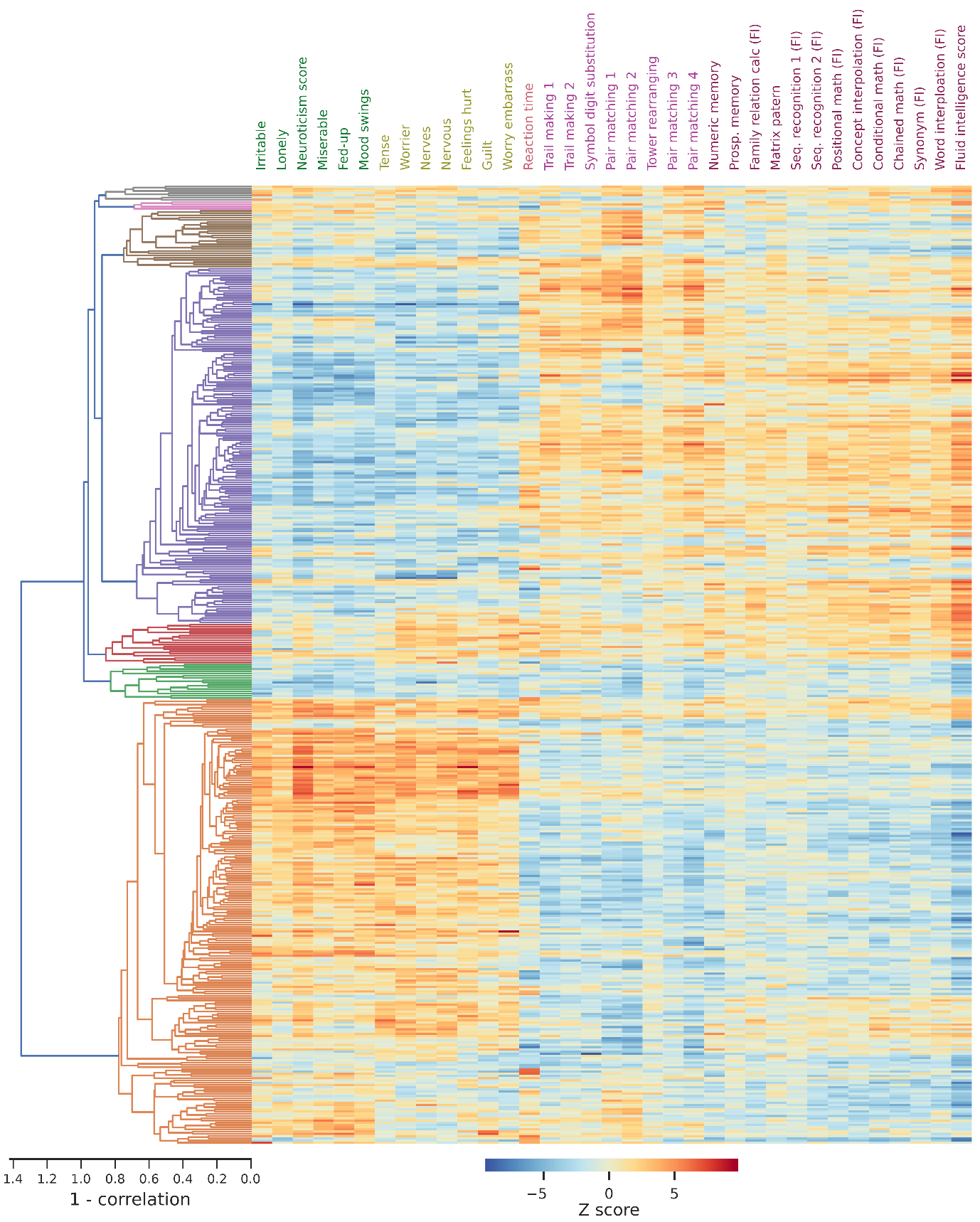

**Supplementary figure 4: Hierarchical clustering of univariate z-scores from 431 lead variants identified by MOSTest multivariate analysis of 35 measures of neuroticism and cognition.**

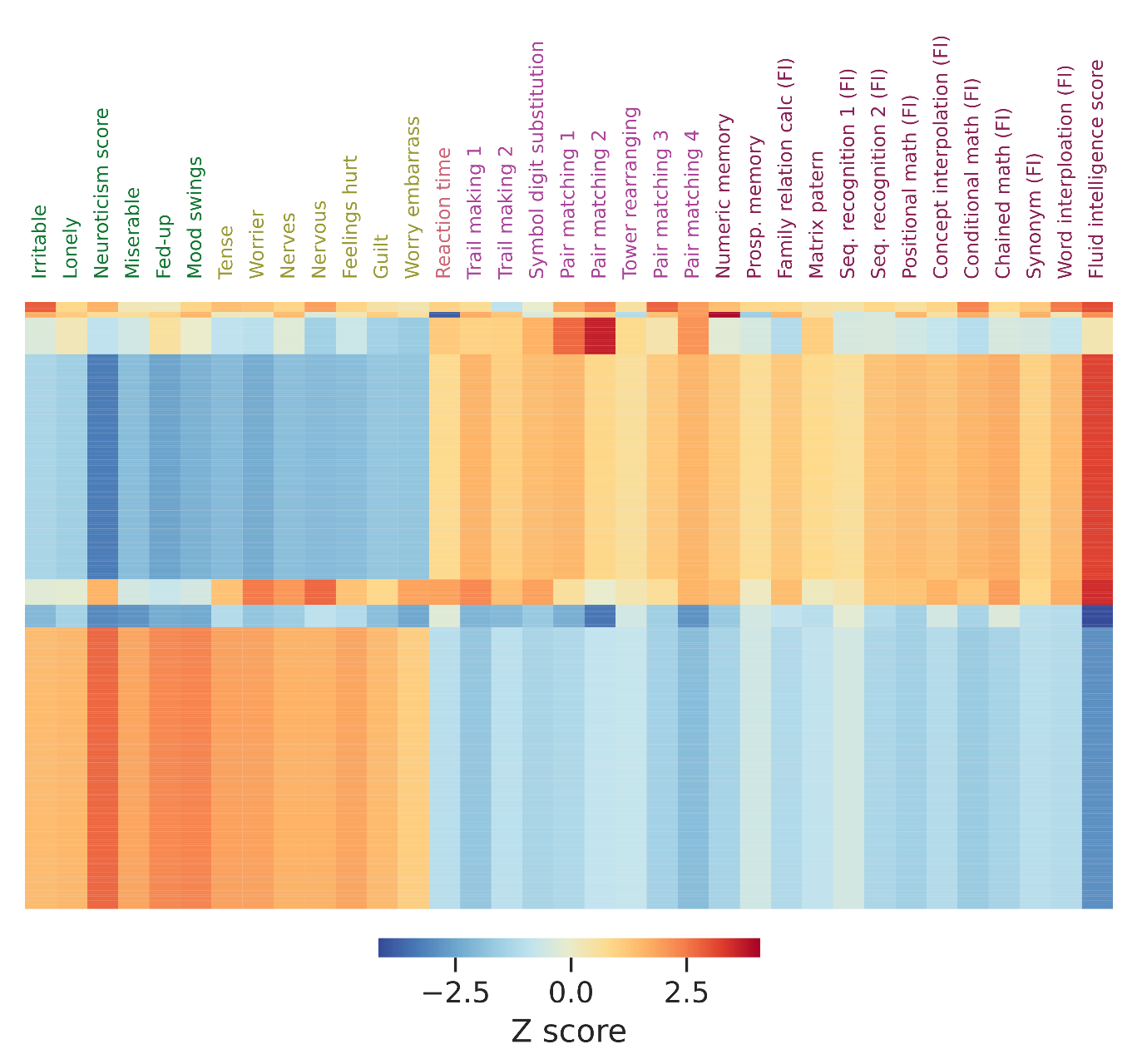

**Supplementary figure 5: Patterns of effect directions across 431 lead variants associated with 35 measures of neuroticism and cognition.** Median value for each of 7 clusters based on hierarchical clustering of univariate GWAS z-scores (supplementary figure 4).

**Supplementary figure 6: Patterns of distributed genetic associations at the SNP-level**. P-values from univariate GWAS of 35 mental traits plotted for the top 40 lead variants, locus number, which corresponds to MOSTest significance rank*,* is provided in brackets. Univariate p-values are plotted on the logarithmic scale as the distance from the centre of each circle plot. Genome-wide significance (p<5x10^-8^) is represented by the dashed line. Positive effect direction is illustrated by a filled circle and negative effect direction by a clear circle. Phenotype clusters are derived from genetic correlation-based hierarchical clustering (**figure2)**

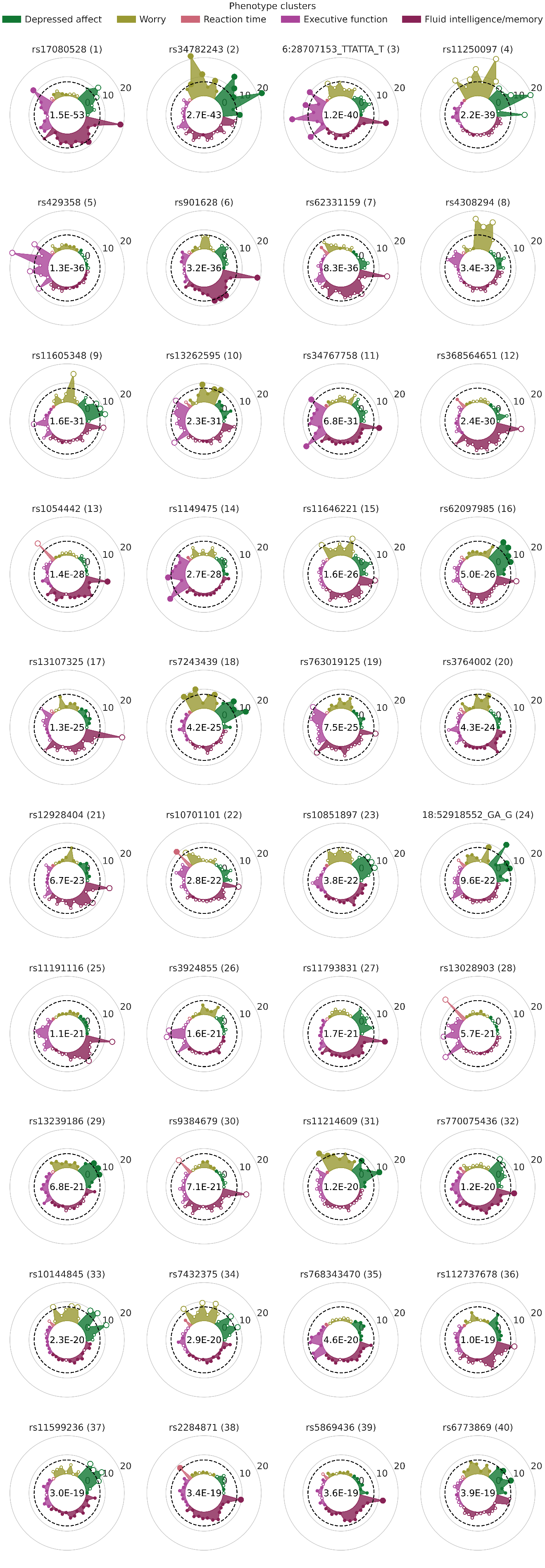

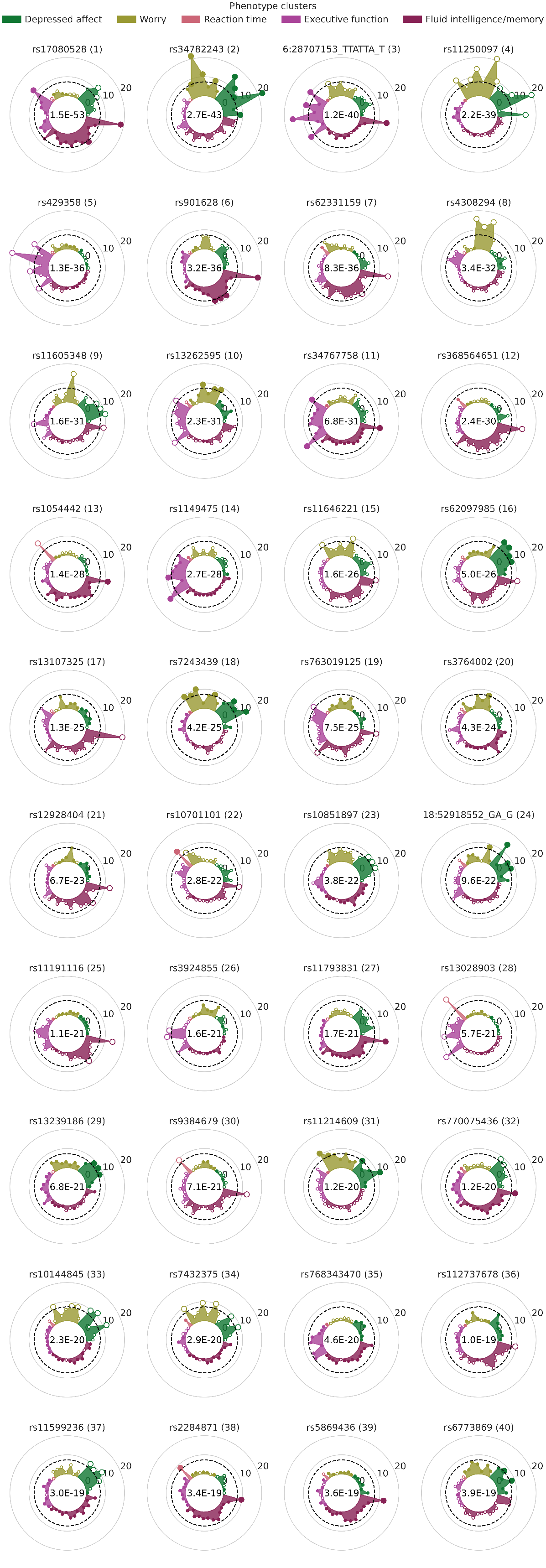

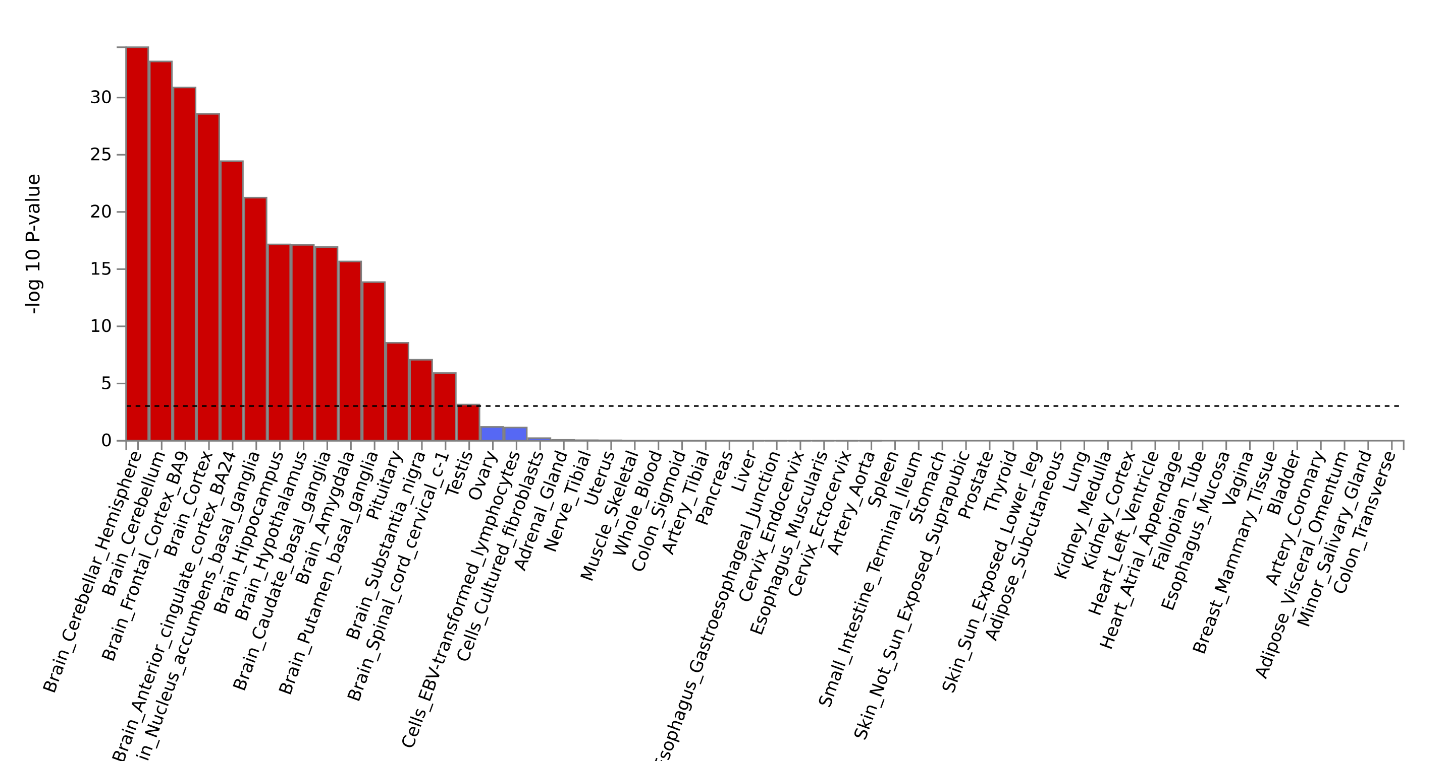

**Supplementary figure 7. MAGMA Tissue expression analysis.** Tissue specific gene expression profiles were tested for their association with multivariate distribution of neuroticism and cognitive traits. This demonstrated high specificity for brain tissues, alongside significant association with testicular tissue.

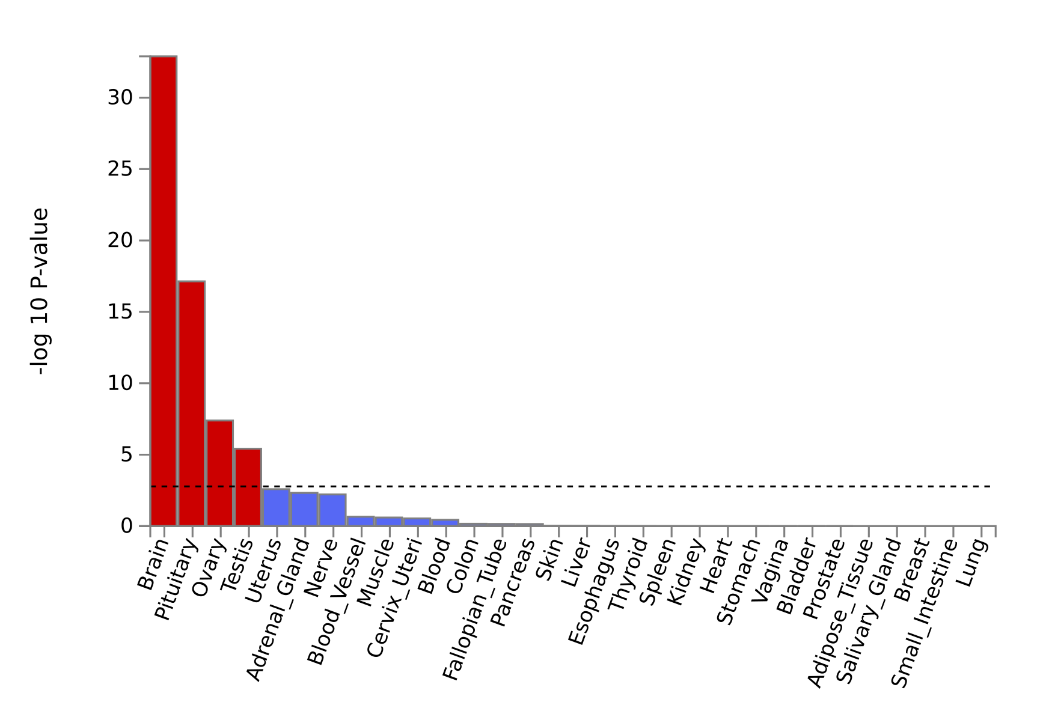

**Supplementary Figure 8. MAGMA Tissue expression analysis – organ level.** Organ specific gene expression profiles were tested for their association with multivariate distribution of neuroticism and cognitive traits. This demonstrated significant association with brain, pituitary, ovary and testis specific genes.

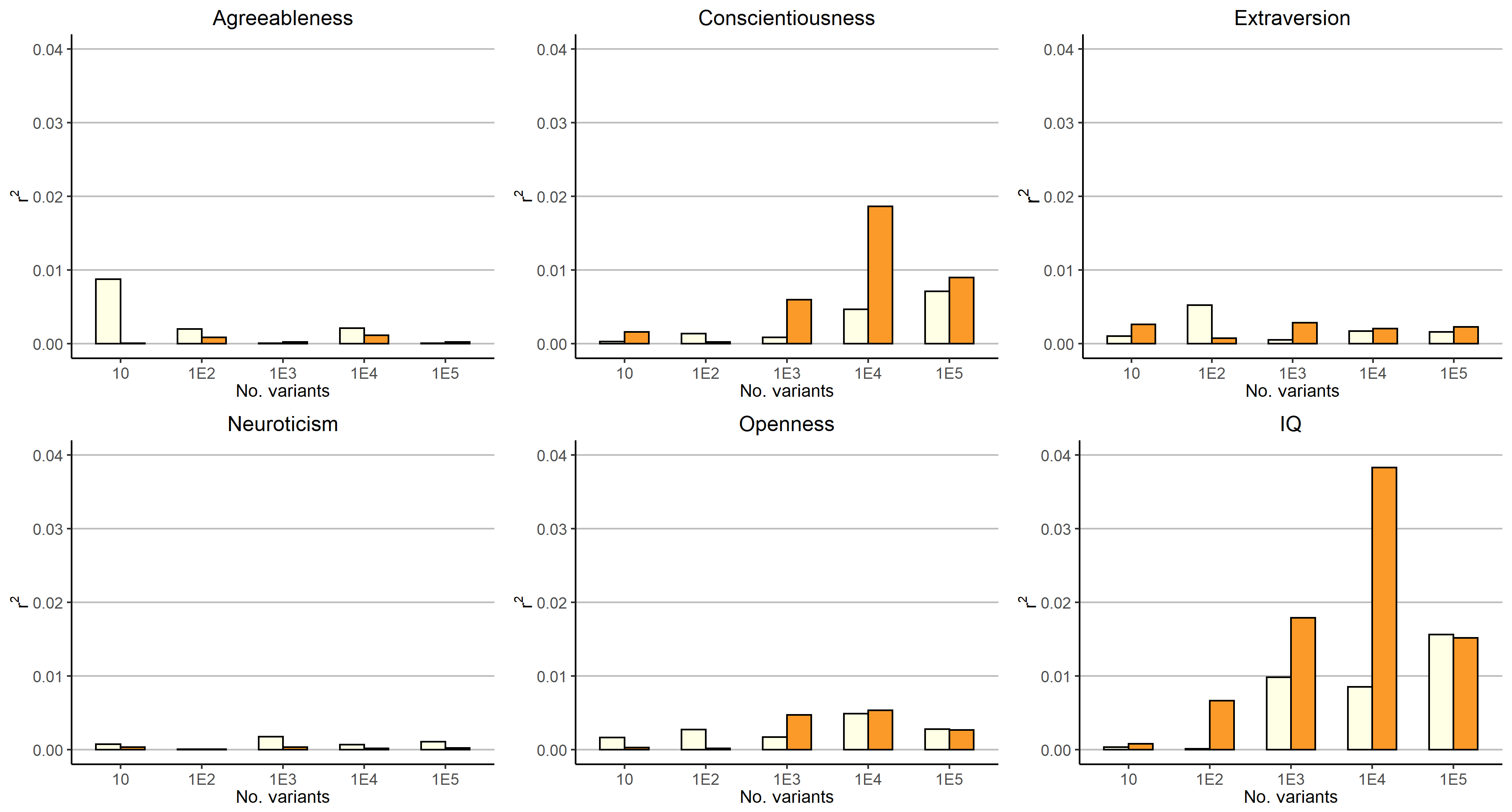

**Supplementary figure 9.** Explained variance of polygenic risk scores (PRS) (r2, y-axis) for top 10, 100, 1000, 10,000 and 100,000 independent variants using primary GWAS-based ranking (i.e. p-values, light grey) and cFDR-based ranking (dark grey) from 23andMe (big 5 personality types) and CHARGE (cognition)GWAS summary statistics. PRSs were tested in healthy participants from the Thematically Organised Psychosis study (n=578-1066).
